# Local homeostasis preserves global neural dynamics compensating for structural loss during human lifespan aging

**DOI:** 10.1101/2025.08.08.669360

**Authors:** Suman Saha, Priyanka Chakraborty, Amit Naskar, Dipanjan Roy, Arpan Banerjee

**Affiliations:** Cognitive Brain Dynamics Lab, National Brain Research Centre, NH48, Gurgaon, 122052, Haryana, India; School of Electronics Engineering, Vellore Institute of Technology, Chennai Campus, Kelambakkam-Vandalur Road, Chennai, 600127, Tamil Nadu, India; Department of Mathematics, Rampurhat College, Dackbanglow More, Rampurhat, 731224, West Bengal, India; School of AIDE, Center for Brain Research and Applications, Indian Institute of Technology, NH 62, Nagaur Road, Jodhpur, 342037, Rajasthan, India

**Keywords:** Aging, anatomical topology, functional networks, metastability, homeostasis, neurotransmitters

## Abstract

Aging brain undergoes a structural decline over lifespan accompanied by changes in neurotransmitter levels, leading to altered functional markers. Past studies have reported human resting state brain display a remarkable preservation of coordination among neural assemblies stemming from an underlying neuro-computational principles along aging trajectories, however, the true nature of which remains unknown. Here, we identify the computational mechanisms with which neurotransmitters, such as altered GABA and glutamate concentrations, can preserve functional integration across lifespan aging, despite structural decline. We employ multiscale, biophysically grounded modeling, constrained by the empirically derived anatomical connectome of the human brain, where the neurotransmitter concentrations can be free parameters that are algorithmically adjusted to maintain regional homeostasis and optimal working point. The two estimated neurotransmitters can maintain critical firing rates in the brain region and mimic age-associated functional connectivity patterns, consistent with empirical observations. We identified invariant GABA and reduced glutamate as the principle computational mechanism that can explain the topological variation of functional connectivity along lifespan, validated using graph-theoretic metrics. The results are subsequently replicated on three distinct datasets. Thus, the study offers an operational framework that integrates brain network dynamics at macroscopic and molecular scales, to gain insight into age-associated neural disorders.

## Introduction

Aging effects are observed at multiple scales of brain organization, ranging from microscopic level alteration in neurotransmitter levels [1–7], to degradation in regional gray matter volume [8], and long-range white matter tracts among brain areas [9–11]. At the macroscopic level, changes in neuronal coordination cause cognitive and behavioral impairments [12–14], or changes to the functional integration in unfolding brain networks [15–17]. Because of such multi-scale interactions, the effects of aging on neural activity become complex. For instance, some functions of the healthy aging brain deteriorate largely, such as declined processing speed, inhibitory function, long-term memory, and working memory [18]. In contrast, other cognitive functions, such as, implicit memory and knowledge storage, remain relatively preserved and show greater resilience to the effects of brain aging [16, 18–22]. Earlier investigations have shown that despite age-associated structural decreases, functional imaging studies reveal somewhat surprising yet pervasive increases in pre-frontal activation with age, reflecting the brain’s adaptive and resilient nature. This helps protect cognitive function during healthy aging, as explained by the scaffolding theory of aging and cognition [18]. Thus, an overarching question arises: can the age-related changes captured by neuroimaging markers be integrated into an operational framework that encompasses the myriad of complex interactions spanning multiple scales of brain organization, including large-scale neural field dynamics and neurotransmitter kinetics?

To conceptualize an operational framework for understanding age-related changes in the brain, one can delve into the structural and functional markers across the lifespan [23, 24]. Typically, graph properties can quantify alterations in brain networks, e.g., modularity and global efficiency [14, 25–27]. For instance, using a longitudinal design, Coelho and colleagues observed that the modularity of whole-brain structural connectivity (SC) increases with age in a moderate-sized group of human participants (N=51) [10, 27]. This general trend also persisted in cross-sectional data from different publicly available cohorts of large sizes (N=174) from three independent datasets used in our study. An increasing number of studies suggest the effective use of graph properties to capture alterations in resting-state functional connectivity (FC) across lifespan aging [16, 23, 28]. Thus, graph-theoretic measures of network segregation, integration and resilience can serve as excellent validation tools to compare age-related variations and reorganization in structural and functional brain networks across cohorts.

In parallel, complexity of neural coordination in time-varying functional networks [29–32] can also provide validation measures to quantify age-related network changes across cohorts. The brain-wide dynamics, involve transitions between “order” - when network nodes may display in-sync activity and “disorder” when nodes are weakly synchronized or display incoherent dynamical relationship [33, 34]. Such transitory “in-between” dynamical state of the brain is assumed to be metastable, which does not commit to a specific attractor state, but remains perpetually prepared to internalize a stimulus that drives the transitions from one attractor state to another [35]. In characterizing global dynamics of the coordinated neural activity at rest, the measure of metastability has been extensively used across various research studies [11, 16, 29, 36]. Being in a metastable state allows the brain to maximize information processing and to promptly react to any internal or external stimulus [37–39]. This dynamical state is often conceptualized as the optimal working point of a healthy brain [30]. Emergence of such neural dynamics depends on two cardinal factors: (i) mutual influences among brain-wide neural populations mediated by their interconnecting anatomical projections (i.e., SCs), and (ii) neurotransmitters responsible for regional firing activity of excitatory and inhibitory populations. While the interconnected fibers serve as the structural foundation for synaptic communication across distributed cortical networks, the neurotransmitter kinetics regulate local excitation-inhibition homeostasis, together enabling the integration of neural activity that supports brain functions and cognitive processes.

Available evidence suggests that qualitative variations of SC properties play an important role in reshaping the brain dynamics differently, likely through their influence on the local ratio, or balance of excitatory and inhibitory cell activity [40, 41]. An earlier study showed how the interplay between structural topology and key controlling parameters, such as excitatory-inhibitory (E-I) balance, played a pivotal role in determining the critical dynamics [42]. An example in the macaque brain showed that a biophysically-grounded computational model, simulated on its connectome, revealed both feedback projections and a heterogeneous distribution of fiber strengths among brain regions as fundamental for the emergence of ignition-like dynamics [43]. These emergent dynamics are thought to be a necessary condition for conscious processing [44]. Ideally, one could rely on the biophysically plausible nonlinear model of neural dynamics to accurately illustrate the importance of anatomical connectivity in supporting coordinated brain dynamics. For example, when simulated by constraining human interregional SC—derived from diffusion-weighted magnetic resonance imaging (MRI), the model replicates empirically observed patterns of interregional FC [34]. Using such a computational framework, numerous studies have demonstrated how the aging brain gradually tunes its intrinsic parameters, e.g., global coupling strength or local E-I regulations [11, 24, 45–48]. This is identified as an effective mechanism that compensates for white-matter loss and upholds its operational working point at maximal metastability. However, the computational principles unifying age-related changes in structural network properties and compensatory mechanisms involving two primary neurotransmitters, GABA and glutamate, have received less attention. Under this background, we hypothesize that healthy aging, where fluid intelligence, language, and other higher-order cognitive functions are typically preserved, must also underlie an invariance of the optimal dynamic working point, indexed by metastability, mediated by the multi-scale interactions involving neurotransmitter kinetics and E-I membrane currents at the neural field level.

To achieve this aim, we adopt the perspective of dynamical compensation underlying healthy aging. We seek to explore mechanistic insights of the adaptive nature of the brain by conceptualizing metastable dynamics as the healthy brain’s optimal working point. The analytical framework integrates the balanced homeostasis regulated by GABA-glutamate with empirical observations on SC and FC graph properties. Accordingly, we employ a biophysically inspired multiscale dynamical mean field (MDMF) model, constrained by human anatomical connectomes. Under optimal parameters, coupling strength, and excitatory and inhibitory neurotransmitters (glutamate and GABA), the model is capable of reproducing resting-state neural activity [46, 49]. The large-scale brain dynamics are modeled by local inhibitory and excitatory neuronal populations with uniform properties across all brain regions. The assumption is based on a homogeneous distribution of the two model parameters, GABA and glutamate concentrations, which are estimated through state-of-the-art model inversion methods. The two free parameters are algorithmically adjusted while preserving desired metastable dynamics within the aging cohorts and achieve a uniform firing rate of approximately 3*Hz* across all brain regions (in line with experimental data [50–52]). Age-related changes in SC properties solely influence the parameter estimation, further validated through qualitative comparison between the empirical observations on FC changes and model-based FC properties. Our model simulation demonstrates that in response to the structural insults, the aging brain engages in a continuous functional reorganization primarily driven by the excitatory neurotransmitter, glutamate, playing a central role in this compensatory process.

## Methods

Fig. 1 depicts multiscale brain organization, and workflow for model simulation and evaluation. Using two biologically inspired constraints, geared towards fitting the spatio-temporal characteristics, we undertake a model inversion to predict variations in GABA and glutamate concentrations mediated by the SC changes with age. Ideally, the static FC provides a temporal averaged snapshot of all covariations pairwise between brain areas, can be considered as a summary measure of spatial features. Thereby, the closeness between simulated and empirical FC is minimized to estimate the two model parameters. Meanwhile, the resting-state brain dynamics shows a tendency to remain at a maximally metastable state [29], quantified by the dynamical measure of metastability that depicts variance in the Kuramoto order parameter [53] fluctuations over time. To match the resting state activity, the measure of metastability is maximized on the two parameter plane, set as another optimization condition. Next, both the conditions are unified to estimate the model based GABA-glutamate values. Finally, we need a set of independent metrics which are not directly related to the method of parameter estimation but can be an independent validation measure for the usability of the model. To meet this end, topological properties of empirical FC computed using graph theoretic metrics can be compared against measures of integration/segregation applied to simulated FC.

**Figure 1.**
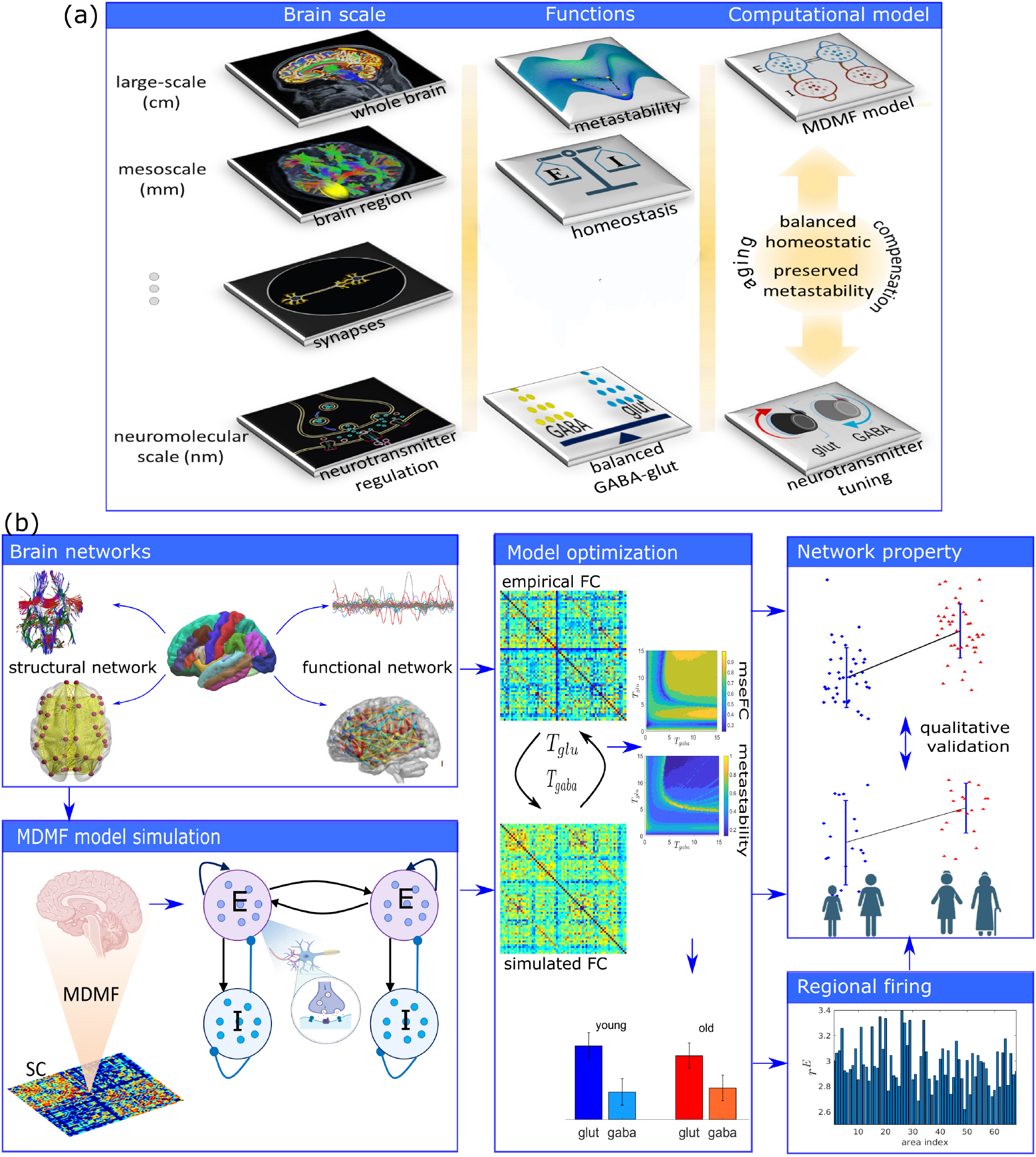
Multiscale brain organization, and workflow for model simulation and evaluation. (a) First column displays a schematic diagram of different brain scales from microscopic to macroscopic level. Second column shows functions at different brain scales. Global dynamics quantified by metastability and E-I homeostasis are observed at macroscopic and mesoscopic scales, whereas, glutamate-GABA kinetics can be observed at microscopic level. Third column presents the working principles of the computational construct. The MDMF model placed at each node of an anatomical brain network (top tile) are optimally tuned by the two parameters (glutamate, GABA) across lifespan aging while maintaining E-I balance. (b) Overview of the entire pipeline for model fitting and evaluation. Empirical structural connectivity (SC) is constrained to the MDMF model that generates simulated FCs. Two conditions, i.e., maximizing metastability and minimizing FC distance between empirical and simulated FCs, are used to estimate optimal Glutamate and GABA values of for individual participants. Based on the optimal parameter values, simulated FCs are generated for individual participants, while the critical firing rates are maintained. The graph theoretic metrics of segregation and integration from the simulated FCs are measured and compared against the empirical FC properties, which provides an independent qualitative validation for the robustness of our results.

### Empirical data description

Data sets for this study were obtained from three different sources. We acknowledge Berlin Centre for Advanced Imaging, Charité University Medicine, Berlin [54] for data collection and sharing. We used the preprocessed data provided by the group. This dataset was collected by a 3T Siemens Tim Trio scanner with a 32-channel head coil (2mm isotropic voxel size). The pre-processed data from the Nathan Kline Institute (NKI) Rockland was obtained from the publicly accessible UCLA Multimodal Connectivity Database (UMCD) [55]. We acknowledge the Cambridge Centre for aging and Neuroscience (CamCAN), University of Cambridge, UK, for data collection and sharing. They used 3T Siemens Tim trio scanner with a 32-channel head coil (voxel size 3×3×4.4mm [56]). We have pre-processed randomly selected participants from similar age groups to CamCAN, Berlin and NKI datasets. Details of participants, data acquisition, and pre-processing are mentioned in the Supplemental Materials with a summary of demographics in Table 1.

**Table 1.** Demographic data across two different stages of adulthood in men and women.

| Data set | Group | Young<br>Number/mean±sd | Old<br>Number/mean±sd |
| --- | --- | --- | --- |
| CamCAN |  | Age range 18-34 | Age range 60-85 |
| | Female | 18 / 26.28 ± 4.43 | 18 / 68.1 ± 6.67 |
| | Male | 19 / 25.57 ± 5.17 | 14 / 70.16 ± 8.08 |
| | Total | 37 / 25.97 ± 4.7 | 32 / 69.14 ± 7.41 |
| NKI |  | Age range 18-30 | Age range 60-85 |
| | Female | 19 / 22.58 ± 2.85 | 16 / 69.69 ± 7.92 |
| | Male | 29 / 22.72 ± 2.81 | 11 / 70.27 ± 9.3 |
| | Total | 39 / 23.08 ± 3.12 | 27 / 70.29 ± 7.62 |
| Berlin |  | Age range 18-33 | Age range 55-80 |
| | Female | 13 / 26 ± 4.73 | 11 / 64.13 ± 7.74 |
| | Male | 12 / 25.33 ± 3.2 | 3 / 58.75 ± 10.4 |
| | Total | 25 / 25.68 ± 4 | 14 / 62.3 ± 8.71 |

### Dynamical measure of metastability

Recent computational brain network models [30, 57] have predicted (for instance) that the human brain operates at maximum metastability during the resting state. It is manifested as an optimal information processing capability and switching behavior [30, 58–60]. Also, the existing cognitive aging theories are explained better with the concept of metastability [16, 29, 60] and used to explore changes in the physiological substrate over aging [16, 61]. We have chosen metastability as one of the optimal conditions to connect the parametric role to the functional organization. Metastability indexes the perpetual state of transition observed in brain signals without committing to a specific attractor state [62–64] and has been used as a metric to capture the dynamic working point of the brain at rest by several researchers [29, 30, 49]. There are several advantages of using metastability such as, the metric can be used to quantify the tendencies of neural systems to operate at segregative and integrative modes [64], characterize the state of criticality where any dynamical system remains on the verge of phase transition thus enhancing information processing capabilities [30, 65]. Finally, the phenomena can be studied via generative models of neural dynamics, and hence allows the researchers to gain mechanistic insights into network mechanisms generating brain signals [49, 66, 67]. Motivated from these knowledge, we have utilized the dynamical measure off metastability to capture the brain dynamics, further we set these as one of the key constraints in our computational framework to optimize the model-based parameters.

First, the Kuramoto order parameter [53], *r*(*t*) is calculated by, 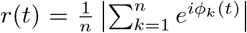. where, *ϕ*_*k*_(*t*) is the instantaneous phase of *k*^*th*^ brain region. The value of *r*(*t*) lies between 0 ≤*r*(*t*) 1; ≤*r*(*t*) = 1 representing complete synchronous, *r*(*t*) = 0 for desynchrony, and (0 *< r*(*t*) *<* 1) representing partial synchrony among all brain regions. Hence, *r*(*t*) indexes the Global coherence among a system of oscillating brain regions. We measure the temporal variability of *r*(*t*) to capture the fluctuations in global spatial synchrony within the whole brain over time. Variation in *r*(*t*) indicates the fluctuation among collective states (such as coherent or incoherent) over time, which is indexed by metastability. Metastability [29, 49, 60] is used to capture the temporal fluctuation, estimating the tendency of a brain region to deviate from the coherent manifold. Thus, it could be a better measurable unit for observing spatial cohesion for a short period (maybe a few hours) and may also serve as a key constituent for observing a gradual change in lifespan brain neural dynamics on a parametric plane (e.g., GABA, glutamate plane) over a very slow time scale, such as the lifespan. Metastability (**M**) can be calculated by taking the standard deviation of *r*(*t*) over time:

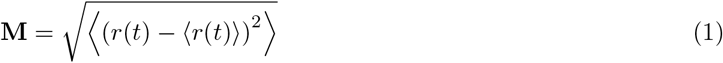

where⟨.⟩ denotes average over time. When compared across data sets, the measure of metastability can extract information about dynamic complexity associated with healthy lifespan ageing (Fig. 2).

**Figure 2.**
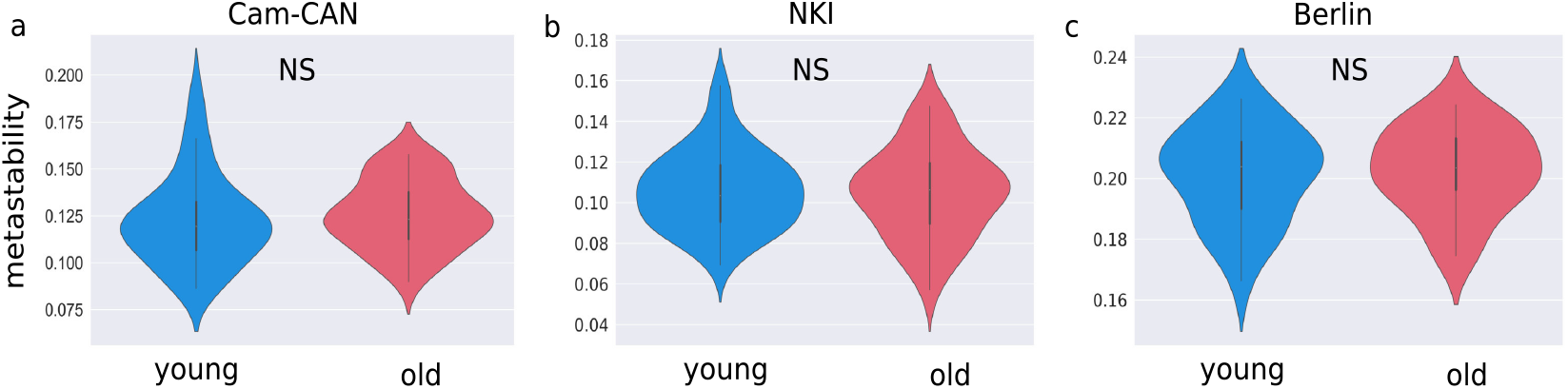
Brain-wide dynamics is quantified by metastability, derived from resting state BOLD activity. Metastability is computed from the empirical BOLD signals of CamCAN (young n=32, old n=37), NKI (young n=39, old n=27), and Berlin (young n=25, old n=14) datasets, display no statistical significance (*p>*0.05) between the young and older adults.

### Preparing empirical SCs and FCs for network analysis

To obtain graph properties, we use undirected-weighted and binary network measures from the Brain Connectivity Toolbox (BCT) (http://www.brain-connectivity-toolbox.net). The structural connectivity (SC) is weighted graph, normalized by the maximum count of white-matter fibers, thus it ranges between 0≤SC≤1 for individual subjects obtained from diffusion weighted imaging (DWI). We derive participant-wise undirected-weighted network properties of the normalized SC. The functional connectivity (FC) is a symmetric matrix that captures the brain-wide long-term correlations between any pair of brain regions, measured by Pearson correlation coefficient ranging between -1≤FC≤1. We convert the original weighted FC matrix into a undirected binary graph by applying a threshold (*δ*) based on the p-values obtained from pair-wise Pearson correlations while taking the pairwise correlation of the BOLD signals between any pair of brain regions. We average the thresholds over entire cohorts to obtain a single threshold value. Thereafter, the weighted FCs are converted into the binary network by setting the condition as, FC_*i*,*j*_=1, if abs(FC_*i*,*j*_) *> δ*, where *δ*=0.076; otherwise, FC_*i*,*j*_=0. In other words, only the significant correlations are chosen to build the binary FC network. Next, we have derived the FC properties for various other threshold values in order to check threshold dependency on the graph theoretical metrics. In particular, the FC thresholding is defined by the minimum correlation coefficient value corresponding to the p-values *<* 0.04 and 0.03, i.e., abs(FC)≥*δ*=0.085 and 0.091. For the three chosen thresholds, preserved edge is between 88% to 85% of the total network’s edge (please see, Supplementary Material Fig. S3). Finally, we tested statistically significant changes in the network properties between two age groups for both brain networks (SC and FC) using independent t-tests and Mann-Whitney U-test analyses for the three data sets Fig. 3 (a-f).

**Figure 3.**
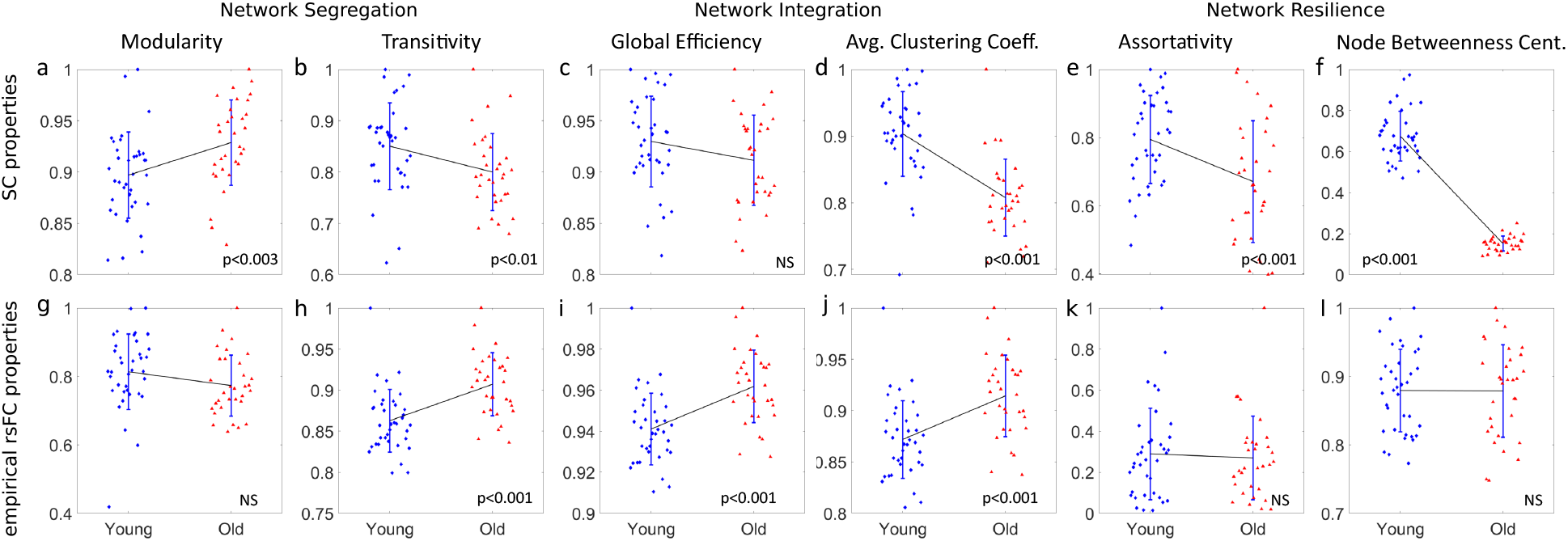
Graph theoretical metrics derived from brain connectomes, empirical SCs and FCs. (a-f) Top row presents the SC properties in two age groups (n=37 young and n=32 older adults). (a) Structural modularity is increased, and (b) transitivity shows a decreasing trend. It can be interpreted as declined structural segregation. Due to loss in white-matter tracts in the aging brain, the connections across modules become more sparse. Thus, segregated SC resulted in a decreased inter-regional exchange of excitation. Though (c) global efficiency remains invariant, (d) significantly decreased average characteristic path length indicates a decline in network integration, which has a direct impact on the network resilience, confirmed by (e) assortativity, and (f) node betweenness centrality comparing the two age groups. (g-l) Functional network properties are shown in the second row (n=37 youmg and n=32 older adults). (g) Functional modularity remains unaltered with age, implying an average number of functional modules remains intact. However, specific brain regions could vary within a module in the two age groups. A significant increase (*p <* 0.001) is observed in (i) global efficiency and (j) average characteristic path length. Increased global efficiency and average characteristic path length in the elderly group signify an increased functional network integration. No significant changes are seen in (k) assortativity and (l) node betweenness centrality between the two groups, which describe unaltered functional resilience in aging brain. To visualize alteration patterns between the age groups, we join the two means by a black line, where the standard deviations from the mean are shown in blue lines. The independent t-test analysis determines significant changes (no changes). Network properties are derived using Brain Connectivity Toolbox [26].

### Definitions of graph theoretical metrics

The network properties were computed using BCT [26]. Brain network’s segregation is captured by measuring modularity and transitivity. Integrative capabilities of a network can be captured by global efficiency and average characteristic path. Structural and functional network resilience or vulnerability is assessed using assortativity and node betweenness centrality. The definition and description of the graph theoretical metrics are:

i. Modularity [25] of a network is 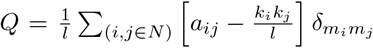, where *m*_*i*_ is the module containing node *i*, and 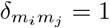, if *m*_*i*_=*m*_*j*_, and 0 otherwise. *a* is a binary adjacency matrix. *l* is the number of links and *N* is total nodes. For a weighted graph, modularity is 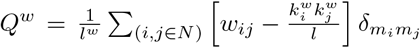, where *l*^*w*^ is the sum of weights in the network; *w*_*ij*_ is the connectivity weight between nodes *i* and 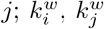 are the weighted degrees of nodes *i* and *j*, respectively. Modularity gives network resilience and adaptability, measuring the degree of segregation. Communities are subgroups of densely interconnected nodes sparsely connected with the rest of the network. In the case of a functional network, modularity signifies coherent clusters of functional modules.
ii. Transitivity [26, 68] is a classic measure of clustering coefficient that captures the tendency of a node to cluster together. High transitivity means the network contains communities or groups of nodes that are densely connected internally. Transitivity of a network is 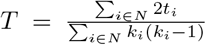, where *k*_*i*_ is the degree of *i*^*th*^-node. Number of triangles around a node 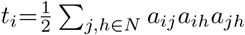. *a* is the adjacency matrix (undirected binary graph). For a weighted graph, geometric mean of triangles around node *i* is 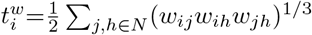.
iii. Global efficiency [69],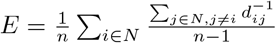, where *N* is the set of all nodes; *n* is the total number of nodes; *d*_*ij*_ is the shortest path length between node *i* and *j*. 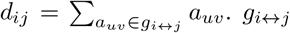 is the shortest path (geodesic). The average inverse shortest path length is the global efficiency, which may significantly contribute to integration in larger and sparser networks [70]. In a weighted graph, the shortest weighted path length between *i* and 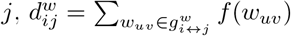, where *f* is a map (inverse) from weight to length.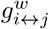 is the shortest weighted path between node *i* and *j*.
iv. Characteristic path length [71], 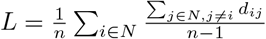, where *N* is the set of all nodes; *n* is the total nodes; *d*_*ij*_ is the shortest path length between *i* and *j*. The characteristic path length for weighted graphs is an estimate of proximity. The global efficiency is the average of the inverse shortest path length. For weighted graphs, we use the shortest weighted path length 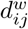.
v. Assortativity [72], 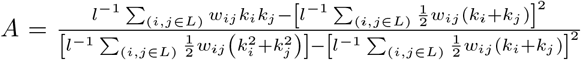, where *k*_*i*_, *k*_*j*_ are the weighted degrees of nodes *i* and *j*, respectively; *w*_*ij*_ is the connectivity weight between nodes *i* and *j*; *L* is the set of all edges within the network, and *l* is the total number of edges. Structural and functional network resilience is assessed utilizing the assortativity [73, 74] and node betweenness centrality [75].
vi. Node betweenness centrality [26, 75] of vertex *i* is 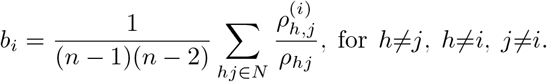 The number of shortest paths between *h* and *j* is given by *ρ*_*hj*_, and 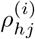 is the number of shortest paths between *h* and *j* that pass through node *i*. Betweenness centrality is computed equivalently on weighted networks, and path lengths are computed on the weighted paths. We take (1*/n*) ∑_*i*∈*n*_ *b*_*i*_ to get the average node betweenness centrality. Betweenness centrality measures the extent to which a vertex lies on paths between other vertices. Vertices with high betweenness have considerable influence within a network under their control over information passing between others. If the vertices are removed, they will cost in disrupted communication, thus depicting the network’s resilience or vulnerability.

### Multi-scale Dynamic Mean Field (MDMF) model

The dynamics of a putative brain area is governed by a stochastic nonlinear dynamical system, the multi-scale dynamic mean field (MDMF) model [49] that describes the homeostatic regulation, the regional excitatory-inhibitory (E-I) neural firing activity. The large-scale brain model is constrained by the SC matrix, and generates simulated FC which is compared against empirical resting state FC to estimate the two free parameters (GABA and Glutamate) in an optimal way. Thereby, the brain-wide coordinated dynamics are emerged from the interplay between brain’s anatomical topology and neurotransmitter levels. Individual brain region is modeled by two distinct pools of excitatory and inhibitory neuronal populations having recurrent E-I, I-I, and I-E interactions, and coupled with neurotransmitters GABA and glutamate via NMDA (N-methyl-D-aspartate) and GABA synapses, respectively. The study by Destexhe and colleague [76] conceptualized the occurrence of neurotransmitter release into the synaptic cleft as a pulse following the arrival of an action potential at the presynaptic terminal. Accordingly, one can consider a chemical kinetic reaction as *R*+*T↔TR*, where T, R, and TR are the neurotransmitters released in the synaptic cleft, unbound receptors, and bound receptors of postsynaptic neuron, respectively. Subsequently, neurotransmitter release can be captured in the rate equation that describes the dynamics of probability of open channels of a specific neurotransmitter (synaptic gating variable). Detailed expressions of how the concentration of GABA and glutamate can be obtained from population-level averaging from individual neuron-level synaptic concentrations are presented in our earlier study [49]. The long-range connections are made between the excitatory pools of any pair of brain regions, defined as SC, which is derived from the fiber densities (interconnected fiber bundles) computed by diffusion tensor imaging data of healthy human adults. The regional excitatory pools receive following input currents: inhibitory currents from local inhibitory pools, self excitatory currents, long-range excitatory currents from the excitatory pools of other brain regions, and a constant external current. The brain-wide dynamics is described by the current-based MDMF model [49] as:

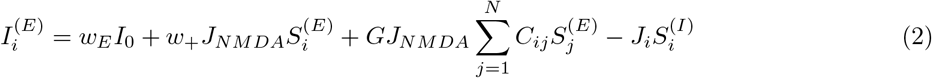

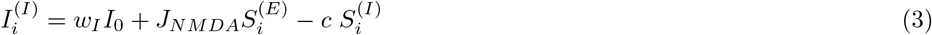

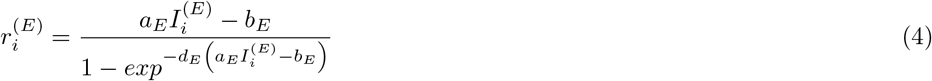

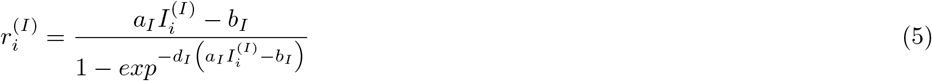

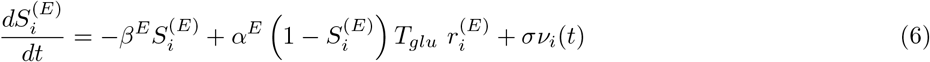

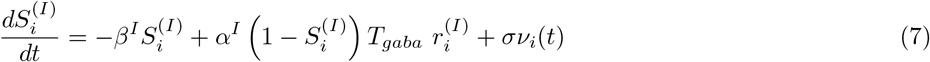

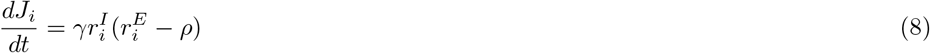

where, the subscripts *i, j* indicate brain areas, *i, j*=1,2,…,*N*. *N* is total number of brain regions and its value is different for different parcellations, e.g., *N* =68 for Desikan-Killiany atlas [77], and *N* =188 for Craddock-200 atlas [78]. The superscripts *E* and *I* represent excitatory and inhibitory populations, respectively. 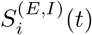 denotes the average excitatory or inhibitory synaptic gating variables of the *i*^*th*^ region. The time-dependent gating variables are drawn by averaging the fraction of open channels of neurons.

The base model proposed in [46] is identical in form for the first four Eqs. (2-5), and the later two Eqs. (6-7) are proposed by Naskar and colleagues [49], by incorporating kinetics of the two primary neurotransmitters, glutamate (*T*_*glu*_) and GABA (*T*_*gaba*_) within the dynamics of the two gating variables 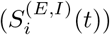. Thus, the MDMF model extends computational observations beyond large-scale neuroimaging, capturing neuromolecular dynamics that govern region-specific E-I balance and neural firing. The two free parameters, *T*_*glu*_ and *T*_*gaba*_, capturing the neurophysiology of the excitatory and inhibitory process, are algorithmically adjusted to fit the model generated FC and empirical FC. For a fixed glutamate and GABA concentration, we select coupling strength *G* for which simulated and empirical rsFC distance is minimum and their correlation is maximum, while maintaining a firing rate approximately 3-4 Hz [79]. The fitting of G and its effects on the parameter estimation is shown in Supplementary Material Fig. S8. Stochasticity is introduced by additive uncorrelated white Gaussian noise *ν*_*i*_ in two gating variables with intensity *σ* for individual regions. Default parameters are given in Supplemental Materials Table S2.

The local synaptic coupling strength from inhibitory to excitatory pools is denoted by *J*_*i*_. Dynamics of the inhibitory feedback is governed by Eq. (8). At the mean-field level, the biological complexity involved in the balance of E-I can be captured grossly using the mathematical implementation of the inhibitory plasticity rule [80]. An inhibitory plasticity rule represents changes in *J*_*i*_(*t*) (synaptic weight) to ensure that the inhibitory current clamps to an excitatory population. Thus, homeostasis is achieved by the dynamics of local inhibitory weights *J*_*i*_(*t*) as a function of time, such that the firing rate of the excitatory population is maintained at the target firing rate *ρ* = 3*Hz*. The chosen target firing rate emerges when E-I balance is achieved. *γ* is the learning rate in *sec*.

We have not considered spatial variability in the neurotransmitters’ concentrations, thus GABA and Glutamate values are homogeneously distributed across all brain regions. Hence, we will get one single pair of parameter values for each independent simulation. For the optimal choice of parameters, the regional excitatory firing rates are approximately 3-4Hz, concurrent with empirical observations [50–52, 79]. Since the model is stochastic in nature, it is unlikely that the converge to a unique solution. Thereby, we have assessed the model performance by comparing simulated FC (generated for the estimated GABA-glutamate values) against empirical FC changes with age. The optimal GABA and Glutamate levels can be estimated from a model inversion approach where the whole-brain dynamics is constrained according to the following conditions: (i) while a normal healthy brain displays metastability at rest, it is maximized on the two parameter plane, and (ii) the mean-square root-error between empirical and simulated FCs (mseFC) is minimized. The concept is elaborated with detailed descriptions and figures; see Fig. S4 in Supplemental Material.

### Simulated BOLD activity

We calculate model generated resting state BOLD signals from the model-based neural activity of the excitatory pool firing rate 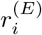 in individuals brain region *i*^*th*^ using the hemodynamic Balloon-Windkassel model in [81, 82]. In the hemodynamic model, an increase in the firing rate 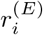 increases vasodilatory signal *s*_*i*_ subjected to autoregulatory feedback. The blood flow *f*_*i*_ in *i*^*th*^ brain region responds in proportion to this signal with concurrent changes in blood volume *v*_*i*_ and deoxyhemoglobin content *q*_*i*_.

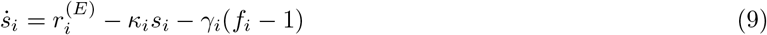

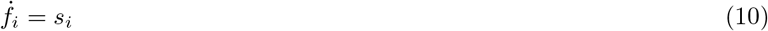

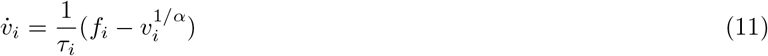

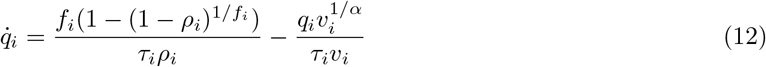

*ρ*=0.34 is the resting oxygen extraction fraction, *τ* =0.98 is a time constant, and *α*=0.32 is the Grubb’s exponent, represents the resistance of the veins. *κ*=0.65 and *γ*=0.41. The BOLD signal for *i*^*th*^ brain region is considered as a static nonlinear function of volume *v*_*i*_ and deoxyhemoglobin *q*_*i*_ comprised of volume-weighted sum of extra- and intravascular signals, given by

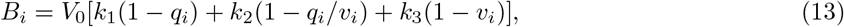

where *k*_1_ = 7*ρ*_*i*_, *k*_2_=2, *k*_3_=2*ρ*_*i*_ − 0.2, and *V*_0_=0.2 is resting blood volume fraction. All the biophysical parameters are taken from [81]. To consider functionally relevant frequency range for resting-state conditions, the simulated BOLD activity has been filtered using a bandpass filter (0.001 Hz*< f <*0.1 Hz). Thereafter, we detrended the simulated BOLD activity, and used Pearson correlation between any pair of brain regions to generate the simulated or model-based FCs.

### Functional connectivity distance

To compare the simulated FC with empirical FC, we calculated the linear distance for each combination of *T*_*gaba*_, and *T*_*glu*_ parameters. The FC distance for a total of *n* regions is calculated using mean-square root-error (mse) between the two FCs as: Please note, to calculate the linear distance between the empirical and

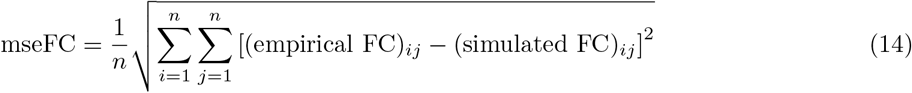

simulated FCs, the weighted FCs are considered, whereas the binarized FCs have been used to derive graph properties.

### Model-based parameter estimation based on spatio-temporal properties

Model inversion is a key step for parameter optimization of complex nonlinear models [83]. The following steps are followed to extract the GABA and Glutamate on an individual participant basis: (a) Structural connectivity matrices are derived from diffusion tensor imaging data of each participant. Detailed descriptions of data acquisition and structural connectivity preparations are explained in Sec. 4 and Supplemental Material Section 1. (b) MDMF model units are placed for each node of the structural connectivity matrix, so that simulated neural activity, e.g., firing rate of excitatory/inhibitory populations at each brain area can be generated. (c) The hemodynamic BOLD activity is then estimated by the Balloon-Windkessel algorithm, to estimate the brain dynamics at each parcel. (d) Pearson correlation is derived from the simulated BOLD time series among brain regions and referred to as simulated FC or model-FC. Simultaneously, metastability is also computed using steps outlined in Sec. 4. (e) The parameter space of GABA/Glutamate (*T*_*gaba*_, *T*_*glu*_) is scouted using two constraint equations addressing the issues of spatial and temporal complexity respectively. Euclidean distance between simulated and empirical FC is minimized, and the parameter distribution, where metastability is maximum, has been identified (Fig. 4(A-E)). The test parameter space for glutamate (*T*_*glu*_) [0.2, 15] and GABA (*T*_*gaba*_) [0.2, 7] are motivated from empirical observations [49] varied along a set of 75×35=2625 discrete, equispaced points with concentrations in units of mMol. For each tuple (*T*_*glu*_, *T*_*gaba*_), we generate simulated BOLD signals and model-based FC for a given SC from one participant which we compare to the empirical FC. As a result, total number of model simulation is 456750 (75×35×174) for the considered cohort of 174 subjects from three datasets. For each subjects total simulation time was set to 7 mins (total 7×60×1000, with 0.1 msec time interval resulting in 4200001 iterations, where first 2 mins are discarded as a transient time). One subject required approximately 22 hours of simulation time on a CPU with 4 cores for parallel computations. Thus, the optimal set of 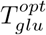 and 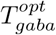 can be conceptualized to be a simple average of the optimal values, separately obtained from spatial and temporal constraint conditions, following the relationship, where *a*=0.5, because, equal weighting is assigned to the measures of spatial and

**Figure 4.**
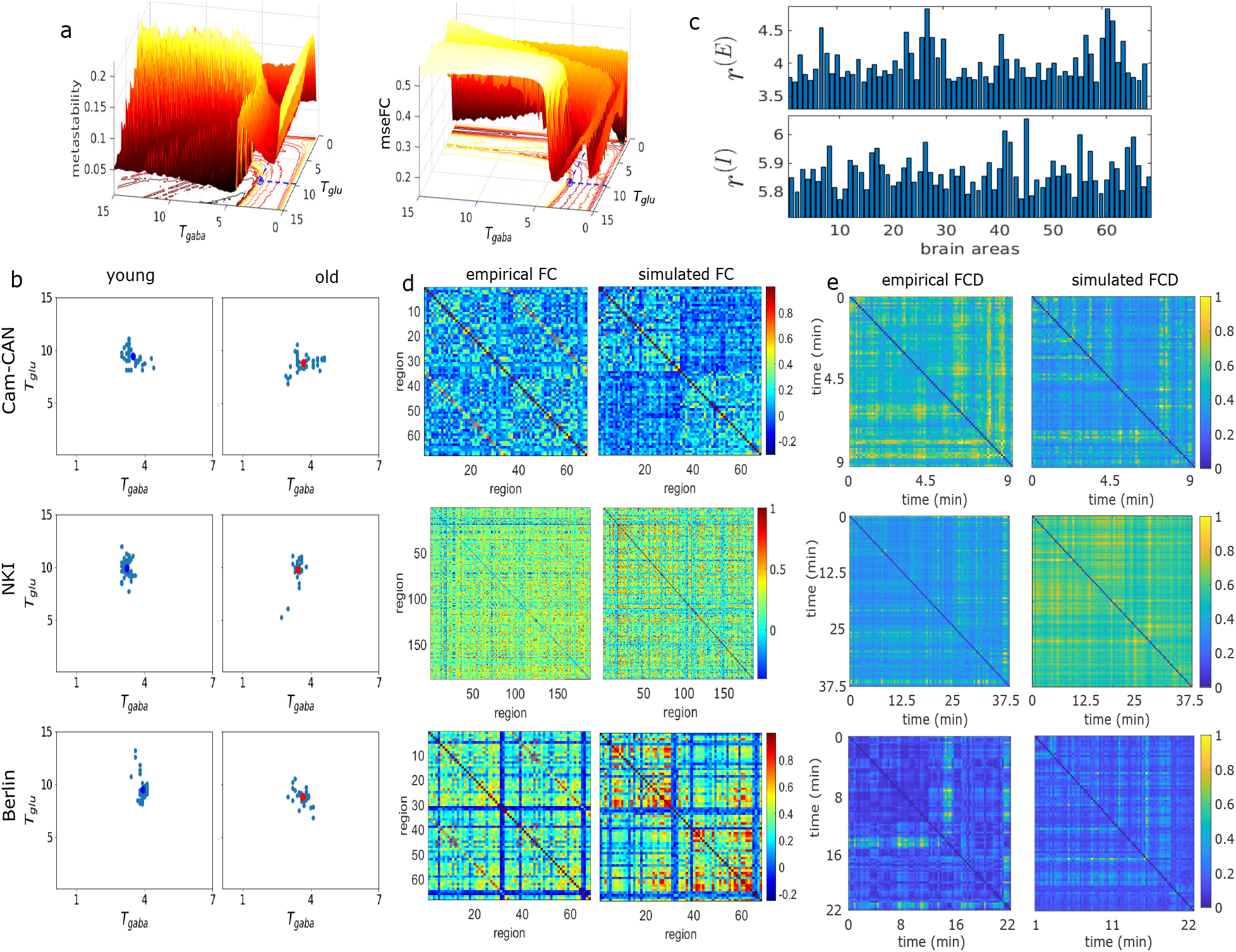
Model-based results obtained from averaged subjects: parameter estimations, firing and FCs. First, the SCs of individual groups are averaged (young n=1, old n=1). Next, 50 independent simulations are performed. (a) The 3-D maps of metastability and mean-square root-error between FCs (mseFC) are plotted on *T*_*gaba*_ *− T*_*glu*_ parameter plane. The blue stars, marked on the 2D parameter plane, indicate optimal *T*_*glu*_, and *T*_*gaba*_ values. (b) Clouds of estimated parameters are shown for young adults and elders, produced by 50 independent simulations. Blue and red circles represent the average values in young and old groups, respectively. (c) Regional excitatory and inhibitory firing rates are plotted for the estimated average *T*_*glu*_, and *T*_*gaba*_ values. (d) Empirical and simulated static FCs are plotted using estimated parameters, taking average over trials. (e) Empirical and simulated FC dynamics (FCD) are shown in time-versus-time plot. All the simulated results are produced using estimated parameter values.

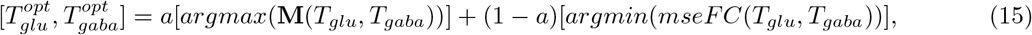

temporal complexity. This way, we obtain each pair of optimal concentrations, *GGC* = (*T*_*glu*_, *T*_*gaba*_)^*opt*^ in mMol. We calculate the GABA-glutamate ratio (GGR) by taking their ratio as *GGR* = (*T*_*glu*_*/T*_*gaba*_)^*opt*^ for individual subjects. Participant-wise estimation steps of glutamate, GABA values using the MDMF model are presented in Supplemental Material Fig. S6. Finally, we perform a two-sample t-test to find statistical differences in the estimated *GGC* and *GGR* between the two age groups in the three datasets.

### Statistics and Reproducibility: Model performance evaluation and test-retest validation

To check the reproducibility of the model-based estimation of GABA-Glutamate values from averaged subjects, we have conducted 50 independent simulations. Clouds of estimated values are shown in Fig. 4, where blue and red circles represent the average values in young and older adults, respectively. Furthermore, to evaluate the generative power of the MDMF model we separated the individual groups in CamCAN and NKI into two cohorts, approximately 40%, labeled as training cohorts (*N*_*young*_ = 15, *N*_*old*_ = 10 for NKI, *N*_*young*_ = 18, *N*_*old*_ = 14 for CamCAN), and 60% testing (*N*_*young*_ = 23, *N*_*old*_ = 17 for NKI, *N*_*young*_ = 24, *N*_*old*_ = 22 for CamCAN) without any overlapping individuals. The training data was used for subject-by-subject estimation of 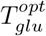 and 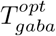 where the input parameter was the SC of each subject. The mean of 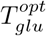 and 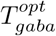 computed for young and elderly groups for CamCAN and NKI cohorts is considered a representative estimate of optimal GABA and Glutamate for each age group.

Simulated FC from BOLD time series, generated using MDMF model, were computed in the testing cohort where the input parameters were the SC of each subject and optimal GABA-glutamate for each age group. Subsequently, the graph theoretic metrics were applied on empirical SC and simulated FC of the test cohort and compared qualitatively with graph theoretic metrics for the entire cohort to evaluate model performance. Model performance is illustrated by measuring graph properties of the model generated FCs for two datasets, shown in model performance evaluation section. Further, we verified our observations for different thresholds utilized to binarize the FCs; please refer to Supplementary Material Fig. S7. The graph theoretical metrics are calculated from empirical FCs using various thresholds (used to binarize the empirical rsFC), which are also used to check the model-generated FCs properties. Finally, to check the robustness of our observation, participant-wise GABA-Glutamate values are estimated for different coupling strengths (*G*), presented in Supplemental Material Fig. S8. To compare between two age groups, we have used independent t-test, or Mann Whitney U-test (for less participants).

## Results

Results are separated into three parts. First, we present the outcomes of graph theoretic network analysis conducted on non-invasive neuroimaging data encompassing both structural and functional aspects derived from diffusion MRI and functional-MRI data. We account for the altered and unchanged graph theoretical metrics in young and elderly cohorts. The empirical observations were validated in three independent datasets. For clear and interpretable comparisons, we purposefully choose two extreme age groups (young adults (18-34) and elders (60-85)). This group-wise design allowed us to capture broad age-related differences in brain network dynamics at the two aging boundaries, and allows us to examine subtle changes occurring in glutamate and GABA levels over a very slow time scale (lifespan aging) that compensates for structural loss, eventually sustaining essential neural dynamics and critical firing rates. To maximize contrasts in glutamate-GABA values and their ratio, we focus on the two opposing ends of the aging spectrum, providing a straightforward observational window to test any contrast in neurotransmitter levels. Our second step of analysis is to elucidate the sensitivity of the parameter estimation process – a key step to effective modelling of empirical FC in different cohorts. We averaged SCs of all the participants within each group. To check trial-wise variability in parameter estimations, we conducted independent simulations on the two averaged SCs corresponding to the two groups, while keeping all other settings fixed. Next, we examine age-related dynamical compensatory mechanisms via computational model dependent prediction of shifts in neurotransmitter to preserve desired brain dynamics, contributing to balance homeostasis, as evidenced from the excitatory-inhibitory firing rates. Finally, we evaluate our model performance, making a qualitative comparison between graph properties of the empirical and simulated FCs.

### Dynamic working point: empirical metastability

To understand changes in the neural coordination dynamics among brain areas across the lifespan, we utilize the dynamical measure of metastability, defined in Eq. (1). Figs. 2(a-c) present violin plots of metastability, derived from participant-wise empirical resting-state BOLD signals for the two age groups. Violin plots of metastability of young and elderly subjects are shown in Figs. 2(a,b,c) for CamCAN, NKI and Berlin data. First, we performed a detrending on the preprocessed BOLD signals of individual brain areas for each subject. Next, the instantaneous phase of each region was determined using the Matlab function ‘hilbert.m’. These two steps are repeated for individual subjects. We checked statistical variations between the two age groups using an independent t-test for CamCAN data and Mann-Whitney U-test for Berlin data. We observed no significant changes in the metastability between the two age groups, suggesting that despite structural changes, the whole-brain dynamical complexity measured by metastability remains unaltered in elderly subjects.

### Alterations in brain networks with age

We have measured weighted network properties of structural connectivity (SC), and for resting-state functional connectivity (rsFC), we have used undirected binary network properties in order to understand the age-dependent alterations in segregation, integration, and resilience features of brain networks. The global graph properties of participant-wise SCs and rsFCs are computed using Brain Connectivity Toolbox (http://www.brain-connectivity-toolbox.net) [26], presented in top row, Figs. 3(a-f), and lower row Figs. 3(g-l), respectively. Statistical significance in the metrics between two age groups is determined by independent t-test (for CamCAN and NKI datasets) and Mann-Whitney U test (for Berlin dataset). Changes in graph properties of SCs and FCs are observed based on segregation [see Figs. 3(a, b) and 3(g, h)], integration [see Figs. 3(c, d) and 3(i, j)] and resilience [see Figs. 3(e, f) and 3(k, l)] of the networks.

### Segregation: Modularity and Transitivity

Segregation is a network property, where dense connectivity is observed within the sub-modules of a large network; however, sparse connections among the sub-modules exist [25, 26]. To characterize network segregation, we computed modularity and transitivity (classical measure of clustering coefficient) of structural and functional networks, shown in Figs. 3(a, b) and 3(g, h), respectively. Modularity signifies how the network is densely connected among the areas physically within a sub-cluster but has sparse interactions across the segregated sub-clusters. A network’s transitivity measures how nodes cluster together. Lower transitivity means the network contains sparsely connected dominant sub-modules. Thus, both measures qualify as network segregation measures.

Increased modularity [14] [shown in Fig. 3(a)] and decreased transitivity [in Fig. 3(b)] along lifespan were observed while applying these measures on structural connectivity matrices. Earlier studies have shown that segregated structure is an outcome of the factors affected mainly by age, e.g., white-matter fiber tracts reductions [10, 24, 84, 85] and degradation in long-range interareal connections [86], thus this is a reconfirmation of previous results across 3 datasets.

Next, modularity and transitivity were applied in resting state functional connectivity (rsFC) to characterize functional segregation. Invariance in modularity [Fig. 3(g)] computed from rsFC implies no change in the number of functional modules or clusters evolved from the resting state BOLD activity in the large-scale brain network. It is observed that the transitivity of structural network was decreased [Fig. 3(b)] in elder subjects, whereas transitivity of functional network increased with age [Fig. 3(h)]. Significantly changed transitivity of FCs in elderly subjects indicates rewiring in the functional connections while preserving modularity, i.e., keeping functional modules intact. An aging brain might calibrate controlling parameters to maintain a self-sustaining pattern in brain dynamics preserving functional modules to avoid functional decline due to altered structure. However, the regions within individual functional clusters or modules may differ across subjects and age groups over time.

### Integration: Global efficiency and characteristic path length

Structural integration signifies that paths are sequences of distinct areas and links. It represents potential routes of information flow between pairs of brain areas [26]. Characteristic path length and global efficiency are measures of global connectedness, providing an estimate of how easily information can be integrated across the network [87]. Global efficiency, related to characteristic path length, is the average of the inverse of shortest path length. Compared to characteristic path length, global efficiency is less influenced by nodes that are relatively isolated from the network [26, 69]. Network integration is observed using global efficiency and characteristic path length from SCs and FCs, shown in Figs. 3(c, d) and 3(i, j), respectively. Structural integration [26] declines with age as suggested by a decreasing trend in global efficiency [14] and significantly decreased characteristic path length [Figs. 3(c, d)] in elderly subjects. In general, the characteristic path length is primarily influenced by long paths (infinitely long paths are an illustrative extreme), while the short paths primarily influence global efficiency [26]. On more extensive and sparer networks, paths between disconnected nodes are defined to have infinite length, and correspondingly zero efficiency, thus affecting long- and short-range information flows across the whole brain.

Functional integration in the brain can be interpreted as the ability to combine specialized information from distributed brain areas rapidly [26]. Paths in binarized functional networks represent an overall projection of sequences of statistical associations and may not correspond to information flow along anatomical connections. Precisely, a functional network’s global efficiency measures a network’s ability to transmit information at the global level [88], which is the inverse of characteristic path length. Increased functional integration, as observed from an increasing trend in global efficiency and average characteristic path length in older groups [see Figs. 3(i, j)], might be associated with the compensatory process against the decline in structural integration [Figs. 3(c, d)], which enhances the communicability or the ability to combine technical information from distributed cortical circuits. The relationship between structure and functions may not necessarily be one-to-one linear mapping, as intrinsic biological parameters largely influence the emergence of collective cortical dynamics.

### Resilience: Assortativity and betweenness centrality

Resilience refers to the ability of a brain network to recover and preserve functions under structural degradation [26]. Some studies have shown that certain brain areas are more prone to overall disruption of function, but several nodes remain unaffected by random perturbations [89]. Networks with fewer influential nodes tend to be more resilient. The commonly used metrics of network resilience are assortativity, and node betweenness centrality [26]. Networks with a positive assortativity coefficient likely have a resilient core of mutually interconnected hubs. In contrast, a negative assortativity coefficient is likely to have widely distributed and consequently vulnerable hubs [26].

Assortativity and node betweenness centrality of SCs and FCs are shown in Figs. 3(e, f) and 3(k, l), respectively. Resilience of the structural network [Figs. 3(e, f)] is significantly decreased in the older group, as observed from a significant decline in assortativity and betweenness centrality in Figs. 3(e) and 3(f), respectively. The assortativity coefficient is a correlation coefficient between the strengths (weighted degrees) of all nodes on two opposite ends of a link. Decreased assortativity indicates a reduced tendency to link nodes with similar strengths. It reconfirmed our previous observations on reduced integration and increased segregation in structure, which could occur due to the loss of connectivity strengths (network became sparse, weighted degrees of individual nodes were decreased) and deterioration in long-range white-matter tracts connecting distant brain regions [24].

On the other hand, resilience of the functional network from rs-FC, measured at the level of individual subjects using assortativity (Fig. 3(k)) and node betweenness centrality (Fig. 3(l)), remains unaltered, essentially no significant changes (*p*=0.6 and *p*=0.45), between two groups. It can be interpreted as a functional compensation among nodes of aging brain against the deteriorated structural resilience (Figs. 3(e, f)), thus upholding functional resilience. We validate our observations on two other data sets, NKI and Berlin data, and the results are presented in Supplemental Materials Fig. S1. We observe similar pattern of alteration and re-organization in empirical SC and rsFC properties, as seen in CamCAN data. Though no statistical significance is seen in the network properties for the Berlin data set (which could be due to comparatively fewer subjects in the two groups), it shows similar alteration patterns as seen in the other two data sets. The results on changes in FC properties are also reproduced for four different thresholds used to generate binarized FC networks, shown in Supplemental Materials Fig. S2 for a curious reader.

### Summary of observations from empirical data

We employed network analysis tools on structural and functional connectivity matrices in young and aged cohorts from three different datasets and report that there are overarching changes in anatomical topology, possibly stemming from displaying changes in white matter fibers, as well as, the measures of segregative and integrative functional network are also altered in aging brain, indicating re-organized FCs. On the other hand, large-scale cortical coordinated dynamics, measured from the resting state empirical BOLD activity, remain preserved with aging, sustaining the desired working point. This led us to hypothesize that a mechanistic approach taken by the aging brain to achieve function preservation may be via changing the neurotransmitter concentrations to maintain a homeostatic equilibrium between excitatory and inhibitory current dynamics. Previous studies have proposed that a homeostatic balance can be maintained when the brain operates at maximal metastability [49]. Thus, the goal of the remaining results section is to identify the age-associated changes in GABA-Glutamate concentrations, using the observation of preserved metastability and aspects of functional networks as proxy measures of constancy of the dynamic working point in a healthy brain [30].

### Model-based observations

#### Simulation results: Optimal parameter selection and consistency

For a discrete grid of 2625 entries, we have generated mseFC and metastability maps for each age group. The 3D maps obtained for mseFC and metastability on GABA-glutamate plane are shown in Fig. 4(a). For each set of (*T*_*glu*_, *T*_*gaba*_), we stored the values of metastability and mseFC measures. The optimal GABA-glutamate values are indicated by blue stars-circles, where the blue dashed lines depict projection on the GABA and glutamate axes. To check the consistency of estimated model parameters, we first averaged the SCs and FCs of the participants from the individual group. To estimate two parameters, we did 50 independent runs with random initial conditions. The scatter plot shows clouds of estimated parameters in Fig. 4(b) for three datasets. The parameters averaged over trials are shown in blue and red circles for younger and older participants, respectively. The optimal parameter set can vary across trials, but remains bounded within a cloud of observation. It may be due to the MDMF model’s high dimensionality, nonlinearity, and stochasticity effects that resulted in different but close estimations for independent runs. For the choice of optimal GABA and glutamate values, the excitatory and inhibitory firing rates remain around 4Hz and 6Hz in all brain regions, shown in Fig. 4(c). The simulated FCs are then generated using the estimated parameter values, placed next to the empirical FCs for the three datasets in Fig. 4(d).

Mahalanobis distance is a pairwise Euclidean distance, calculated between the dominant dynamic FC subspace of one time point and all other time points. The Mahalanobis distances are derived using the algorithms proposed in [32, 90], and the temporal stability matrices are plotted for empirical and simulated FC dynamics (FCD) in Fig. 4(e). These results are produced using the estimated parameter values.

Due to the higher parcellation scheme used in the NKI dataset, the static FC patterns exhibit more complexity and appear lacking in features compared to FC generated from Berlin and CamCAN. However, qualitative similarity in dFC patterns across all data sets can be clearly visible in all three datasets. Overall, simulated FC and dFC when compared for each cohort CamCAN, NKI, or Berlin, using MDMF model at optimal Glutamate-GABA values showed high degree of similarity with empirical data.

#### The MDMF model-based glutamate-GABA concentrations

Estimated glutamate-GABA concentrations (GGC) and GABA/glutamate ratio (GGR) at the level of individual subjects from the two age cohorts are shown in Fig. 5, results from CamCAN data in Fig. 5(a, b), NKI data in Fig. 5(c, d) and Berlin dataset in Fig. 5(e, f).

**Figure 5.**
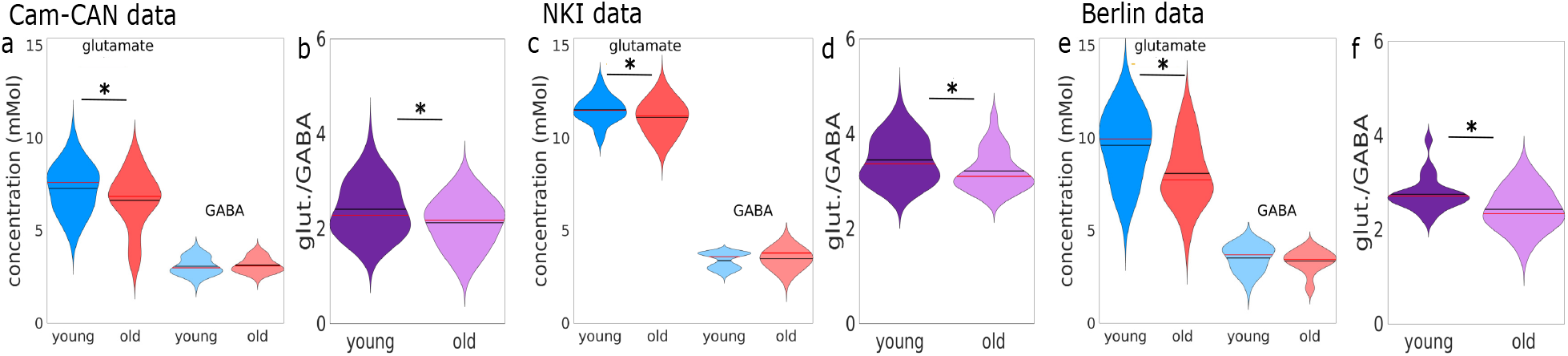
GABA-Glutamate parameter estimation using the MDMF model inversion. Participant-by-participant basis GABA-Glutamate values and their ratio are shown in (a, b) for CamCAN (young n=37, old n=32), (c, d) for NKI (young n=39, old n=27), and (e, f) Berlin data (young n=25, old n=14). The aging brain significantly tunes the glutamate level more prominently than that of GABA level, where a significant decay in GGR can also be observed. The star * indicates *p <* 0.05.

Figure 5(a) shows glutamate and GABA concentrations in young and old groups from CamCAN. Glutamate level is decreased in older groups, where GABA remains almost unchanged. A notable change in the ratio is observed (*p <* 0.05) in Fig. 5(b). The average values of glutamate and GABA for the younger and older group are shown in the black star and green circle, respectively. The scattered blue stars and red circles are the estimated glutamate and GABA values for individual subjects, separating into young and older groups. The three data sets found a similar pattern of altered GABA glutamate levels with age. Further, we have performed a similar analysis on the NKI data set, presented in Fig. 5(c, d). Fig. 5(c) shows glutamate and GABA concentrations in young and old groups. Glutamate level is decreased in older groups, where GABA remains almost unchanged. A notable change in the ratio is observed (*p <* 0.05) in Fig. 5(d). We conducted similar analysis on Berlin data. Figure 5(e) shows violin plots of estimated concentrations of the two metabolites in two age groups, where glutamate level is decreased significantly (*p <* 0.05), but GABA level remains invariant. A significance decrease in GABA-glutamate ratio is observed from the model-estimated results between two age groups, see Fig. 5(f). Furthermore, we illustrate the effect of varying global coupling strength (*G*) on GABA/glutamate estimation in Supplementary Material Fig. S8. This figure shows: (i) optimal GABA/Glutamate estimates at different *G* values, (ii) excitatory and inhibitory firing rates obtained from a single participant, and (iii) empirical and simulated FCs from both a young adult and an elderly participant. Our analysis reveals that there exists a range of *G* values where the simulated FC fits well with the empirical FCs. Most importantly, while the estimated glutamate/GABA values remain consistent across this range, only the excitatory and inhibitory firing rates show variation. The altered level of glutamate concentrations and glutamate-GABA ratio can be interpreted by unaltered neural dynamics, as quantified by the dynamical measure of metastability, which is preserved across aging. Altered glutamate giving rise to the desired complex dynamics essentially reorganized functional connections as evidenced from the empirical network metrics, such as increased transitivity (clustering coefficient), increased integration and invariant resilience in functional network.

The model simulated results suggest that the aging brain alters the optimal operating point [30] by manipulating the two model parameters, in particular by reducing glutamate, to compensate for age-related variability in collective dynamics and avoid functional decline. A study using empirical data has found similar decreasing trends in the concentrations of glutamate in the left hippocampus (HC) and anterior cingulate cortex (ACC) with age [91]. The decreasing trend in glutamate concentration suggests a link between the neurotransmitter and the possible adaptive mechanism of the aging brain to compensate for structural degradation. In our study, we observed that the re-orientation in FC network results from the re-organization in the underlying brain dynamics when the two neurotransmitters control the intrinsic compensatory processes to maintain an equilibrium in the E-I ratio at rest, where the excitatory firing rate sustains at a critical range of 3−4Hz. We must mention that the two neurotransmitters in this study, are only the model parameters that control the dynamics of the MDMF model. We did not attempt to validate the absolute levels of the two neurotransmitters in the context of aging or compare them with the empirical data. All the results on neurotransmitters are theoretical predictions about possible parameter requirements for maintaining desired brain dynamics in a changing structural network.

#### Model performance evaluation: Graph theoretical metrics computed from simulated FC

A test-retest validation is conducted to assess the model based observations. Consequently, our goal is to retest the model performance under estimated optimal parameter values as test condition and empirical results as true cases. In particular, we aim to evaluate the extent to which the model-based FCs, generated using averaged estimated parameters calculated by splitting the cohorts, can replicate similar patterns of changes observed in the empirical FC properties when comparing the two aging cohorts. First, the estimated optimal values of glutamate and GABA from a few subjects (see Methods 2.2.5) are calculated and averaged for each age group. The average values corresponding to the two datasets are: (i) CamCAN: 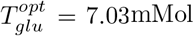 and 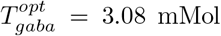 for young; 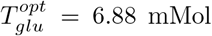 and 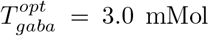 for old; (ii) NKI: 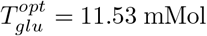 and 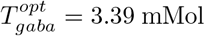 for young and 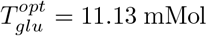 and 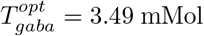 for old. Using these parameters and subject-by-subject SC, we computed the BOLD time series using MDMF (Eqn. (7)) followed by computing FC using Pearson correlation. Next, we binarized the FC and calculated graph theoretical measures at the individual level, shown in Fig. 6, two bottom rows. We generate the binarized FC network for a threshold *δ* = 0.081 corresponding to a *p <* 10^−4^, such that *FC*_*ij*_=1, if *abs*(*FC*_*ij*_) *> δ*; otherwise *FC*_*ij*_=0. To compare the graph metrics estimated across age groups, we have performed the Mann-Whitney U test, the non-parametric alternative to the independent sample t-test due to fewer test subjects. The upper rows display empirical FC properties for CamCAN and NKI data, see Fig. 6, top two rows; two bottom rows show results from the CamCAN and NKI datasets, for simulated FC network properties in blue and red dots for young and older participants, respectively. The threshold for statistical significance is considered to be *p <* 0.05, and is not significant marked by NS. Due to a smaller number of participants in the Berlin data set within two age groups, we did not perform the predictive analysis to evaluate the model performance. We performed test-retest validation of the simulated FC network prediction for two different thresholds used to binarize the simulated FCs, presented in Supplementary Material Fig. S7. Of all measures, only the centrality of the node betweenness is not fully predicted from the test subjects of NKI data, even with varying thresholds (used to build binary FCs).

**Figure 6.**
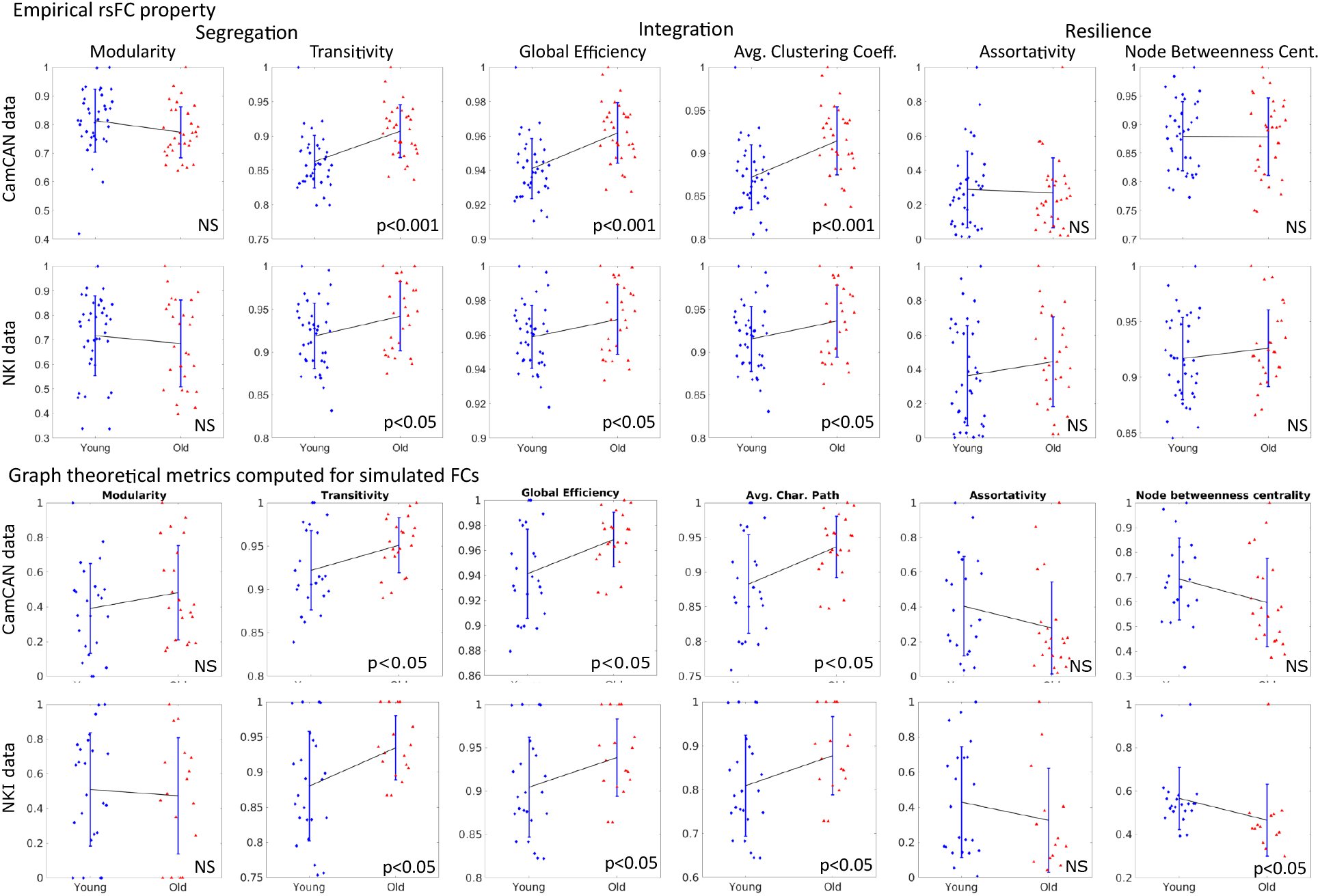
Qualitative comparison between empirical and simulated FC properties. Top two rows display network properties of the empirical rsFC, compared against the observations from simulated FCs (bottom two rows), which are obtained for the optimal parameter choices. Number of test subjects are: n=23 young and n=22 older adults for CamCAN data, and n=23 young and n=17 older adults for NKI data. The variability is coming solely from individual’s SC and is mainly assessed in the test cohort. The MDMF model inversion reproduces almost similar alterations between two age groups occurring for the *test cohorts*, where, the modeling parameters are computed from *training cohorts*. Except, alterations in betweenness centrality is not fully predicted from the test subjects of NKI data. Statistical differences are assessed by Mann-Whitney U test, where *p<*0.05 implies statistically significance, otherwise not significant (NS). A solid black line connecting two mean values of younger and elderly groups, is plotted to visually tracking alterations. The errorbars show standard deviations.

The graph metrics from the MDMF model for both NKI and CamCAN datasets display qualitatively similar variation across age groups, as we observed from the network properties of empirical resting-state FCs between the two groups. These results validate the predictive power of the MDMF model and give us confidence in its use to characterize how the aging brain alters its biological parameters to preserve the collective dynamics around a desired operating point, which may allow the brain to react to any external or internal stimulus flexibly. The alteration can also be interpreted as how the aging brain reorganizes its functional connectivity by re-adjusting the controlling parameters in the face of structural degradation in healthy aging.

## Discussion

Several critical reviews have highlighted the urgent need for tools that can leapfrog studies of neuroscience from a scale-specific biological observation to explain ongoing behavior towards the development of comprehensible multi-scale models that can generate an understanding of the interactions among multiple organizational scales [92–94]. This study focuses on a whole-brain computational model that captures the interactions between two observational levels of neural complexity, ultra-slow BOLD-fMRI dynamics, and neurotransmitter kinetics. The precise interactions transcending the biological scales of observations allow the estimation of critical markers such as Glutamate-GABA concentrations from model inversion techniques across healthy human aging. First, we illustrate that functional segregation, integration, and resilience measures can be preserved and sometimes enhanced across the healthy aging process, even though similar metrics applied on structural network topology point out a gross degradation. Second, we identify a key invariant that captures the dynamic working point of the brain, as well as provides a link to model the interaction between neurotransmitter kinetics and emergent metastable dynamics of the local field with aging. Third, we could estimate the optimal concentration changes in GABA and glutamate associated with healthy aging by model inversion under the constraint of an optimized dynamic working point [30]. Finally, a test-retest validation allowed us to evaluate the predictive capability of MDMF based neurotransmitter estimation. We assessed the model performances under the estimated parameter sets, i.e., assessing the extend to which the model generated results are able to capture similar age-related changing patterns observed in the empirical FC properties. Thus, our study developed a computational microscope based on a biophysically inspired generative neural mass model linking large-scale neuroimaging data with microscopic parameters. The advantage of using this model is two-fold, reconciling observational accounts of metabolite mapping with mechanistic insights of neuronal network workings and possibly setting up future studies that can use this framework to predict the onset of neurodevelopmental and neuroinflammatory disorders where the local E/I balance is crucially perturbed. We discuss these issues in more detail in the remaining sections.

### Structure-function markers of healthy aging

The most salient feature of our empirical network analysis is that the graph theoretical measures, segregation, integration, and resilience, derived from SCs and resting state FCs, showed alterations with age (Fig. 3). Our observations are in line with earlier reports on age-related structural changes [17, 27], and functional reorganization [95–97]. Using multiple network measures such as characteristic path length and global efficiency gives us a set of tools to capture variability in functional network’s integration across subjects [98], between groups [87, 99–102] and are of immense practical importance in brain network study [103, 104], however, cautionary approach must be followed when mechanistic interpretations about engagements of neuronal populations are made, using measures of statistical dependency. In the field of aging research, tracking a gamut of network measures can examine the idea of neurocompensation [105**?**] by which the brain areas improve upon their segregative, integrative and cooperative behavior in the face of structural decline [24, 106, 107]. Being at metastable state, the system is essentially situated in a transient state where it can have more opportunities of choosing an attractor, this can also be interpreted as a state where information processing capabilities increase, leading to higher flexibility [16]. Interestingly, metastability calculated from empirical BOLD signals was unaltered between the young and elderly participants. This can be interpreted as the preservation of information processing capability across the aging cohorts, who, for the most part, are capable of complex cognitive functions (although showing a decrease in performance indices over age, see [31]) emerge from underlying compensatory mechanisms that preserves the global dynamical complexity of the brain. From the perspective of the current study, we use this observation as a constraining tool for model inversion using the MDMF model [49] to estimate the Glutamate and GABA concentration changes across aging trajectories.

### Insight from the model simulation

A key contribution of the present study was the development of a framework that could estimate the local synaptic GABA, Glutamate concentrations (GGC) and GABA, Glutamate ratio (GGR) from non-invasive fMRI. We observed that age negatively correlated with glutamatergic regulation, which was reduced with healthy aging, in concurrence with earlier evidence [13, 108–110]. It is important to note that in the absence of brain-wide MRS data on GABA/glutamate concentrations, evaluating a goodness of fit of model estimated neurotransmitter concentrations is out of scope of the present manuscript. Rather, this work seeks to provide a deeper insight on a probable computational principle by which interaction between brain network dynamics and neurotransmitter kinetics gives rise to a complex adaptive process associated with brain aging. To gain an empirical view of neuromodulation in the aging brain, we refer to the existing literature reporting regional changes in metabolites. For example, empirical observations from healthy human adults in the posterior cingulate cortex [109] and rodent brain [13, 108], revealed significant reduction in the glutamate level in the aging brain, aligned with the patterns of alteration observed in our model-based prediction. The reduced glutamate level has been interpreted as a key mechanism by which the aging brain adapts to maintain the desired regulation of I-E by down-regulating glutamate and also decreasing the expression of several other presynaptic markers [111]. Another study reported a similar negative correlation with glutamate contents in the aging brain in the motor cortex, but a positive correlation with other metabolites, for example, glutamine, N-acetyl aspartate and creatinine [2]. Further, the reduction of glutamate in the motor cortex is linked with the neuronal loss/shrinkage during aging [2]. Other empirical examples showed significant changes at 20% to 50% in the glutamate contents in the frontal cortex, hippocampus, and cerebral cortex with aging [112, 113]. Nevertheless, a review on the critical perspective of aging and development on glutamatergic transmission had categorically shown the effects of aging based on the shreds of evidence from various in-vitro and in-vivo studies on rodent brains, also includes contrasting observations [114]. For instance, studies also reported non-significant changes in glutamate content in the dorsal prefrontal cortex, sulcal prefrontal cortex, temporal cortex, and medial prefrontal cortex during aging [115, 116].

One of the crucial observations that emerge from this theoretical approach is that age-related alterations are different in GABA than glutamate, even though we varied GABA-glutamate concentrations homogeneously across brain areas. The estimated glutamate content was significantly reduced, whereas GABA showed very less variability with age. Similar alteration pattern in the two metabolites was observed an earlier work in the context of aging [110], and also in rodent brain that significantly influenced cortical plasticity [111]. However, in contrast to our observation on GABA, studies have re-ported age-dependent reduction in GABA concentration derived by magnetic resonance spectroscopy (MRS) in human subjects. In particular, reduced GABA was evidenced in the motor, visual, auditory, somatosensory areas, and the perisylvian region of the left hemisphere, which caused an abnormal information processing [3, 4]. Our model simulation showed significant decay in GABA-glutamate ratio in older adults compared to the younger participants, consistent with the MRS-based experimental observation in the PCC/precuneus [117].

Overall, our model simulations suggest that, despite structural decline, the brain-wide neural dynamics are preserved across aging trajectory through readjustments in neurotransmitter levels and their ratio, leading to unaffected E-I balance. Thereafter, the interplay between tuned metabolite levels and cortical plasticity provokes a gradual functional re-organization in the aging brain over a prolonged time (lifespan), while maintaining desired homeostasis. To this point, a critical failure of the present MDMF model is the inability to predict accurate changes in glutamate and GABA levels in individual brain regions. Thus, for future work, it will be advisable to make model-based predictions of heterogeneous distribution of metabolites in the brain which we discuss further in the following sub-section. Nonetheless, much fine-tuning of glutamatergic activity occurs via modulation of the NMDA receptors, which are invisible to MRS because of its limitation in detecting a neurochemical concentration within a localized region of tissues (typically in the order of functional pools of GABA as they might be more tightly bound to macromolecules than others, rendering 2×2×2 cm^3^) [118]. Available evidence suggests that MRS can not separate between different them less “visible” to MRS [119, 120]. In that case, the proposed theoretical framework could be a supporting tool to look into the age-related fluctuations in the metabolites at rest and task-evoked activity, besides MRS-driven data, within a localized area and across whole-brain.

### Limitations and future directions

The computational framework we have presented in this article gives a bird’s eye view of the neuromolecular mechanisms at the whole-brain level. Nonetheless, the biology of the brain is way more complex and our model also has several limitations. First, spatial heterogeneity of the two metabolites in the cortex is not considered in this study. In future, incorporating training models that consist of template MRS data from individual brain regions can be used to address the limitation of regional heterogeneity in the two metabolites. An atlas of whole brain metabolite map from future MRS studies can be used to tune the MDMF model for better predictions of resting state BOLD dynamics that can improve model fits to empirical FC properties. This would also require the implementation of a more advanced model inversion technique beyond two-parameter (GABA, Glutamate concentrations for this manuscript) fitting with optimized metastability and mseFC used here. In other words, machine learning models such as Bayesian model inversion can replace this step. Some other potential limitations are: (i) An earlier observation [121] on a rat brain experiment showed alteration in GABAergic and glutamatergic behavior affected by the age were highly probable to occur in a presynaptic mechanism than in a postsynaptic mechanism. On the other hand, the MDMF model does not distinguish between the pre- or postsynaptic concentration of GABA-glutamate [121, 122]. (ii) Uptake and release of glutamate [113] cannot be separated using the present method. The model does not explicitly tell about the two major sub-types of GABA, i.e., GABA_A_ and GABA_B_. (iii) Measure of total GABA concentration remains unclear whether the GABA signal represents cytoplasmic, vesicular, or free extracellular GABA [118].

It is crucial to note that the standard deviation of instantaneous Kuramoto order parameter as a measure of metastability has a significant limitation: it is only meaningful when applied to rhythmic dynamics. To overcome this limitation, we implemented a narrow band filter (0.001 Hz to 0.1 Hz) on both the empirical and simulated BOLD signals prior to calculating metastability. This filtering process effectively enhanced the rhythmicity of the BOLD signals, thereby ensuring that our metastability measurements remained valid and interpretable throughout our analysis.

While experimental research has demonstrated that metabolite alterations aid in early Alzheimer’s disease (AD) diagnosis [123] and that GABA-mediated excitation-inhibition (E-I) balance restoration can treat depression [124], the in-silico paradigm offers complementary mechanistic insights into disrupted neural dynamics. For instance, computational studies have revealed how noise in the left temporal lobe impairs functional connectivity in AD [125], a framework further extended by recent in-silico perturbation protocols for dynamic recovery [126, 127]. In a long run, the theoretical construct of multiscale modeling can be a useful tool in designing personalized model to identify dynamical mechanisms of imbalanced homeostasis and region-specific vulnerabilities in neural circuit dysfunction across neuropsychiatric disorders and neuropathological conditions Levy and Degnan (2013), e.g., neural migration disorder [128], Parkinson’s disease [129], AD [123], attention deficit hyperactivity disorder [130] and autism [131, 132].

## Acknowledgments

SS was supported by NPDF, SERB-DST, India, File ID: PDF/2021/000585. DR was supported by Ramalingaswami Fellowship, Department of Biotechnology (DBT), India, Award ID: BT/RLF/Re-entry/07/2014 and Department of Science and Technology (DST), India, Award ID: SR/C-SRI/21/2016. AB was supported by award ID: F.NO.K-15015/42/2018/SP-V from The Ministry of Youth Affairs and Sports, Government of India and NBRC Flagship program, award ID: BT/MED-III/NBRC/Flagship/Flagship2019 from the Department of Biotechnology, Government of India. Authors acknowledge the Neuroscience Gateway [133] for facilitating the resources required for model simulation. We acknowledge BioRender for the icons and assets used in Figure 1 (Created in https://BioRender.com). We extend our sincere thanks to the editors and reviewers for their invaluable suggestions and thoughtful comments that greatly helped improve the manuscript.

## Author contribution

SS and AB conceived the study; SS, PC, and AB performed the analysis; AN, DR, and AB contributed tools; SS and AB wrote the first draft; all authors reviewed and edited the manuscript; DR and AB supervised the research.

## Additional information

### Data availability

Data can be downloaded for CamCAN from https://www.CamCAN.org/ and NKI Rockland from https://fcon1000.projects.nitrc.org/indi/pro/nki.html. The Berlin data cannot be shared due to Ethical guidelines of the project. For replication purposes, numerical source data associated with Figs. 2-6 can be found in Supplemental Data 1-5, respectively.

### Code availability

The codes for all the analyses to generate the figures are provided at https://bitbucket.org/cbdl/agingmdmf/src/master/.

**Correspondence** and requests for materials should be addressed to Arpan Banerjee.

## Supplemental Material for

## 1 Descriptions of the three datasets

### 1.1 CamCAN dataset

#### 1.1.1 Subjects

In our study, total 69 healthy participants were included from the Cambridge Centre for Ageing and Neuroscience (CamCAN) cohort http://www.mrc-cbu.cam.ac.uk/datasets/camcan/http://www.mrc-cbu.cam.ac.uk/datasets/camcan/ Shafto et al (2014); Taylor et al (2017).

#### 1.1.2 Data acquisition

Resting state MRI (T1-weighted image), diffusion weighted MRI, and functional MRI were performed using a 3T Siemens TIM Trio scanner with a 32-channel head-coil at Medical Research Council (UK) Cognition and Brain Sciences Unit (MRC-CBSU) in Cambridge, UK.

High-resolution 3D T1-weighted data was collected with a magnetization prepared rapid gradient echo (MPRAGE) sequence using Generalized Auto-calibrating Partially Parallel Acquisition (GRAPPA) with acceleration factor of 2 and other parameters were: repetition time (TR)=2,250 ms, echo time (TE)=2.99 ms, flip angle: 9^°^, field of view (FOV)=256 × 240 × 192 mm; voxel size = 1 mm^3^ isotropic, inversion time (TI)=900 ms, acquisition time (TA)=4:32min Shafto et al (2014).

Diffusion data were acquired in a twice-refocused spin echo sequence with TR of 9100 ms, TE of 105 ms, FOV of 192 × 192 mm, isotropic voxel size of 2 mm^3^, 66 axial slices using 30 directions with b = 1000 s/mm^2^, 30 directions with b = 2000 s/mm^2^, and three b = 0 images with single average Shafto et al (2014). Resting state eye closed fMRI was collected with EPI sequence with 1970 ms TR, 30ms TE, 78^°^ flip angle, 192 × 192 mm FOV, 3 × 3 × 4.44 mm^3^ voxel size, 32 slices of thickness 3.7 mm Shafto et al (2014).

#### 1.1.3 DTI processing

Diffusion MRI data were processed locally using own pre-processing pipeline by using MRtrix3 (Tournier et al (2019), https://github.com/MRtrix3/mrtrix3), FSL http://www.fmrib.ox.ac.uk/fsl http://www.fmrib.ox.ac.uk/fsl, and ANTS http://stnava.github.io/ANTs/, http://stnava.github.io/ANTs/. Main preprocessing steps include denoising (MRtrix command ‘dwidenoise’, Veraart et al (2016)), Gibb’s ringing artefacts removal (MRtrix command ‘mrdegibbs’, Kellner et al (2016)), motion and eddy current corrections (MRtrix command ‘dwifslpreproc’, Andersson et al (2003)), biasness corrections (MRtrix command ‘dwibiascorrecT using ANTs, Tustison et al (2010)). Then a brain mask in DTI space was calculated for each subject using ‘beT command (in FSL) on the preprocessed image. To find the orientation of the fiber(s) (fiber orientation distribution, FOD) in each voxel, we used multi-shell multi-tissue constrained spherical deconvolution (MSMT-CSD; MRtrix command ‘dwi2response’ and ‘dwi2fod’, Dhollander et al (2018)). After that, a global intensity normalization has been performed to make the fiber orientation distributions comparable between subjects. We used Anatomically Constrained Tractography (ACT, MRtrix command ‘5ttgen’, Smith et al (2012)) to generate a tissue-segmented image. In order to use ACT, we first preprocess the T1-weighted image using MRtrix, then we co-registered that image to the DWI using both MRtrix and FSL. We created an mask of the gray-matter/white-matter-boundary, which was useful for streamline seeding. Finally, probabilistic tractography (MRtrix command ‘tckgen’) has been used to generate 20 million tracks. The tractogram was filtered (SIFT2 approach, MRtrix command ‘tcksift2’) further to a find subset of streamlines such that the streamlines densities were more close to fibre densities Smith et al (2015). Each steps were visually assessed and edited by research personnel.

#### 1.1.4 T1 processing

Freesurefer http://surfer.nmr.mgh.harvard.eduhttp://surfer.nmr.mgh.harvard.edu Dale et al (1999) ‘reconall’ used to reconstruct a two-dimensional cortical surface from a three-dimensional volume acquired from T1-weighted image. Recon-all steps were skull stripping from the anatomical image, estimation of interface between the white matter and grey matte, generation of white and pial surfaces. Cortical parcellation was performed using the Desikan-Killiany (DK) atlas Desikan et al (2006), with 68 ROIs.

#### 1.1.5 Structural connectivity

A Desikan–Killiany (DK) atlas based whole-brain connectome was generated for each subjects by computing the fiber density between each pair of ROIs (MRtrix command ‘tck2connectome’ with option ‘scale invnodevol’) to count how many streamlines from one region reach every other regions.

#### 1.1.6 fMRI preprocessing using Desikan–Killiany atlas

Functional MRI (fMRI) images for resting state has been preprocessed using CONN toolbox Whitfield-Gabrieli and Nieto-Castanon (2012) https://web.conn-toolbox.org/https://web.conn-toolbox.org/, a Matlab/SPM-based software. Data were preprocessed using a default preprocessing pipeline of CONN. The preprocessing methods were unwrapping using field-map images, realignment to correct for motion, slice timing correction, segmentation, normalisation to the MNI template, outlier rejection and functional smoothing. Spatial smoothing was performed using a Gaussian kernel with full width of 12 mm. Denoising was used to remove signal changes related to white matter, cerebrospinal fluid, motion, breathing and cardiac pulsations. Finally temporal band-pass filtering at (0.004-0.1 Hz) and linear detrending were performed. For region of interest (ROI) analysis, mean regional BOLD time series were estimated in 68 parcellated brain areas of Desikan–Killiany atlas Desikan et al (2006).

#### 1.1.7 Resting state functional connectivity using Desikan–Killiany atlas

The same participants were subjected to a functional MRI (fMRI) scan during which their eyes-closed awake resting-state data were acquired. The resting-state BOLD activity was recorded for 22 minutes (TR=1.97 sec). The BOLD activity was then down-sampled to fit the 68 ROIs. Aggregated BOLD time series of each region were z-transformed. Next, pairwise Pearson correlation coefficient were computed to obtain 68×68 rsFC matrix for individual subjects.

### 1.2 NKI/Rockland data set

We tested our hypothesis on another human neuroimaging dataset obtained from Nathan Kline Institute (NKI)/Rockland sample (available in the UCLA Multimodal Connectivity Database (UMCD)Brown et al (2012).

#### 1.2.1 Structural connectivity

Diffusion Tensor Imaging (DTI) based structural connectivity matrices and rsFC/Pearson correlation matrix of 75 healthy participants were obtained from the Nathan Kline Institute (NKI), Rockland sample Brown et al (2012). The participants consisted of 30 males and 35 females, with a total age range of 19-85 (mean ± std). We separate 40 participants into two age groups (48 young and 27 old) and averaged their SCs over individual subject for our analysis (see Table 1). Cortical gray matter was parcellated using the Crad-dock 200 atlas Craddock et al (2012). Elements of the SC represented the number of white matter tracts between gray matter parcels. The reader may also refer to the original work Brown et al (2012) for details regarding structural and functional connectome pre-processing. Prior to running the models on the NKI SC, the SC was scaled down by dividing every element by the maximum value found in the original SC consisting of white matter tract numbers between regions. Diffusion tensors were estimated using Diffusion Toolkit (http://trackvis.org/blog/tag/diffusion-toolkit) and tractography was run using the fiber assignment by continuous tracking algorithm Mori and Van Zijl (2002). For each ROI, all fibers were counted that intersected at least one voxel in the source ROI and at least one voxel in any target ROI using custom code, (http://ccn.ucla.edu/wiki/index.php/UCLA Multimodal Connectivity Package). Thus, 188×188 structural connectivity was obtained.

#### 1.2.2 Resting-state functional connectivity using Craddock-200 atlas

Resting state fMRI data was pre-processed using the pipeline described in Brown et al (2012). After marking flagged TRs, the mean time series for each ROI was calculated and then correlated with all remaining ROI time series (excluding flagged TRs) to derive a 188×188 resting-state functional connectivity (rsFC) matrix parcellated using the Craddock-200 atlas Craddock et al (2012).

#### 1.2.3 Resting-state functional connectivity using AAL parcellation

We have performed every analysis in NKI dataset using Craddock atlas; only metastability is measured using the AAL atlas to maintain consistency with the CamCAN analyses, where we also examined the metastability metric using AAL atlas (116 ROIs), besides Desikan–Killiany atlas (see section 2.4.3, Fig. S4 for CamCAN results using AAL atlas). This served as a cross-validation of the metastability metric. Since metastability is a time-varying measure, qualitatively similar results are obtained whether we use Desikan (68 ROIs) or the AAL atlas (116 ROIs).

Functional MRI images for resting state have been preprocessed using the CONN toolbox Whitfield-Gabrieli and Nieto-Castanon (2012) https://web.conn-toolbox.org/https://web.conn-toolbox.org/, a Matlab/SPM-based software. Data were preprocessed using a default preprocessing pipeline of CONN. The preprocessing methods were unwrapping using field-map images, realignment to correct for motion, slice timing correction, segmentation, normalisation to the MNI template, outlier rejection and functional smoothing. Spatial smoothing was performed using a Gaussian kernel with full width of 2 mm, and TR=2.5 sec. Denoising was used to remove signal changes related to white matter, cerebrospinal fluid, motion, breathing and cardiac pulsations. Temporal band-pass filtering at (0.004-0.1 Hz) and linear detrending were performed. Mean regional BOLD time series were estimated in 115 brain areas of AAL atlas Tzourio-Mazoyer et al (2002). We used the preprocessed BOLD signals in NKI dataset for metastability analysis only, presented in the main text Fig. 2.

### 1.3 Berlin data

#### 1.3.1 Subjects

The study included 36 healthy subjects (22 females, 14 males). The written consent from the participants are taken care by Schirner et al (2015). The earlier study Schirner et al (2015) was approved by ethics committee of the Charité University Berlin, and all experiments were performed in compliance with the relevant laws and institutional guidelines. We divided 41 subjects into two age groups comprising 22 young participants ranged in age range 18-33 years (mean age=25.68 ± 4 years, 13 female) and 16 elderly participants of age range 55-80 years (mean age=65.06 ± 7.39 years, 11 female).

#### 1.3.2 Data Acquisition

T1 structural magnetic resonance images (MRI) and diffusion-weighted images (DWI) were acquired at Berlin Center for Advanced Imaging, Charité University Medicine, Berlin, Germany. MRI was performed on a 3T Siemens Trim Trio scanner and a 12 channel Siemens head coil (voxel size). Structural (T1-weighted high-resolution three-dimensional MPRAGE sequence; TR=1900ms, TE=2.52ms, TI=900ms, flip angle=9^°^, field of view (FOV)=256mm×256mm×192mm, 256 × 256 × 192 matrix, 1.0mm isotropic voxel resolution), diffusion-weighted (T2-weighted sequence; TR=7500ms, TE=86ms, FOV = 192mm × 192mm, 96×96 matrix, 61 slices, 2.3mm isotropic voxel resolution, 64 diffusion directions), and fMRI data (2-dimensional T2-weighted gradient echo planar imaging blood oxygen level-dependent contrast sequence; TR=1940ms, TE=30ms, flip angle=78^°^, FOV=192 × 192mm^2^, 3 × 3mm^2^ voxel resolution, 3mm slice thickness, 64 × 64 matrix, 33 slices, 0.51ms echo spacing, 668 TRs, 7 initial images were acquired and discarded to allow magnetization to reach equilibrium; eyes-closed resting-state) were acquired on a 12-channel Siemens 3 Tesla Trio MRI scanner at the Berlin Center for Advanced Neuroimaging, Berlin, Germany.

#### 1.3.3 Structural connectivity

The structural connectivity (SC) for each subject and age is generated using the pipeline described by Schiner et. al. Schirner et al (2015). In the pipeline, high-resolution T1 anatomical images were used to create segmentation and parcellation of cortical and sub-cortical gray matter, white matter segments. The main pre-processing steps for T1 anatomical images involved skull stripping, removal of non-brain tissue, brain mask generation, cortical reconstruction, motion correction, intensity normalization, WM, and subcortical segmentation, cortical tessellation generating GMWM and GM-pia interface surface-triangulation and probabilistic atlas-based cortical and subcortical parcellation. Cortical grey matter parcellation of 34 region of interest (ROI) in each hemisphere was undertaken following Desikan-Killiany parcellation Desikan et al (2006). The connection strength (a value ranging from 0 to 1) between each pair of ROIs was estimated by probabilistic tractography algorithm. SC matrices were generated from each subject’s MRI data and then summed element-wise to obtain an averaged SC matrix. The pre-processing steps for the diffusion MRI data were eddy current and motion correction with re-orientation of b-vectors (b-zero image was linearly registered to the subject’s anatomical T1-weighted image). Tractography was constrained by seed, target, and stop masks. The fiber length was represented in millimeters. The 68 ROIs or nodes had no self-connection loops meaning that the diagonal values of the SC matrix are all zero.

#### 1.3.4 Resting state functional connectivity

The participants were subjected to a functional MRI scan, when eyes-closed awake resting-state data were acquired. The resting-state BOLD activity was recorded for 22 minutes (TR=2 sec) using a 3T Siemens Trim Trio scanner, and a 12 channel Siemens head coil (voxel size). The BOLD signal was computed for 68 ROIs, parcellated by Desikan-Killiany atlas Desikan et al (2006) and z-transformed. Pearson correlation coefficient, between each pair of region, defines resting-state FC matrix.

**Table S1.** Demographic data across two different stages of adulthood in men and women.

| Data set | Group | Young<br>Number/mean±sd | Old<br>Number/mean±sd |
| --- | --- | --- | --- |
| CamCAN |  | Age range 18-34 | Age range 60-85 |
|  | Female | 18 / 26.28 ± 4.43 | 18 / 68.1 ± 6.67 |
|  | Male | 19 / 25.57 ± 5.17 | 14 / 70.16 ± 8.08 |
|  | Total | 37 / 25.97 ± 4.7 | 32 / 69.14 ± 7.41 |
| NKI |  | Age range 18-30 | Age range 60-85 |
|  | Female | 19 / 22.58 ± 2.85 | 16 / 69.69 ± 7.92 |
|  | Male | 29 / 22.72 ± 2.81 | 11 / 70.27 ± 9.3 |
|  | Total | 39 / 23.08 ± 3.12 | 27 / 70.29 ± 7.62 |
| Berlin |  | Age range 18-33 | Age range 55-80 |
|  | Female | 13 / 26 ± 4.73 | 11 / 64.13 ± 7.74 |
|  | Male | 12 / 25.33 ± 3.2 | 3 / 58.75 ± 10.4 |
|  | Total | 25 / 25.68 ± 4 | 14 / 62.3 ± 8.71 |

## 2 Results on the empirical data analysis

### 2.1 Graph theoretical metrics of empirical SC and FC network

We did test-retest on three datasets to check the robustness of our observations.

**Figure S1.**
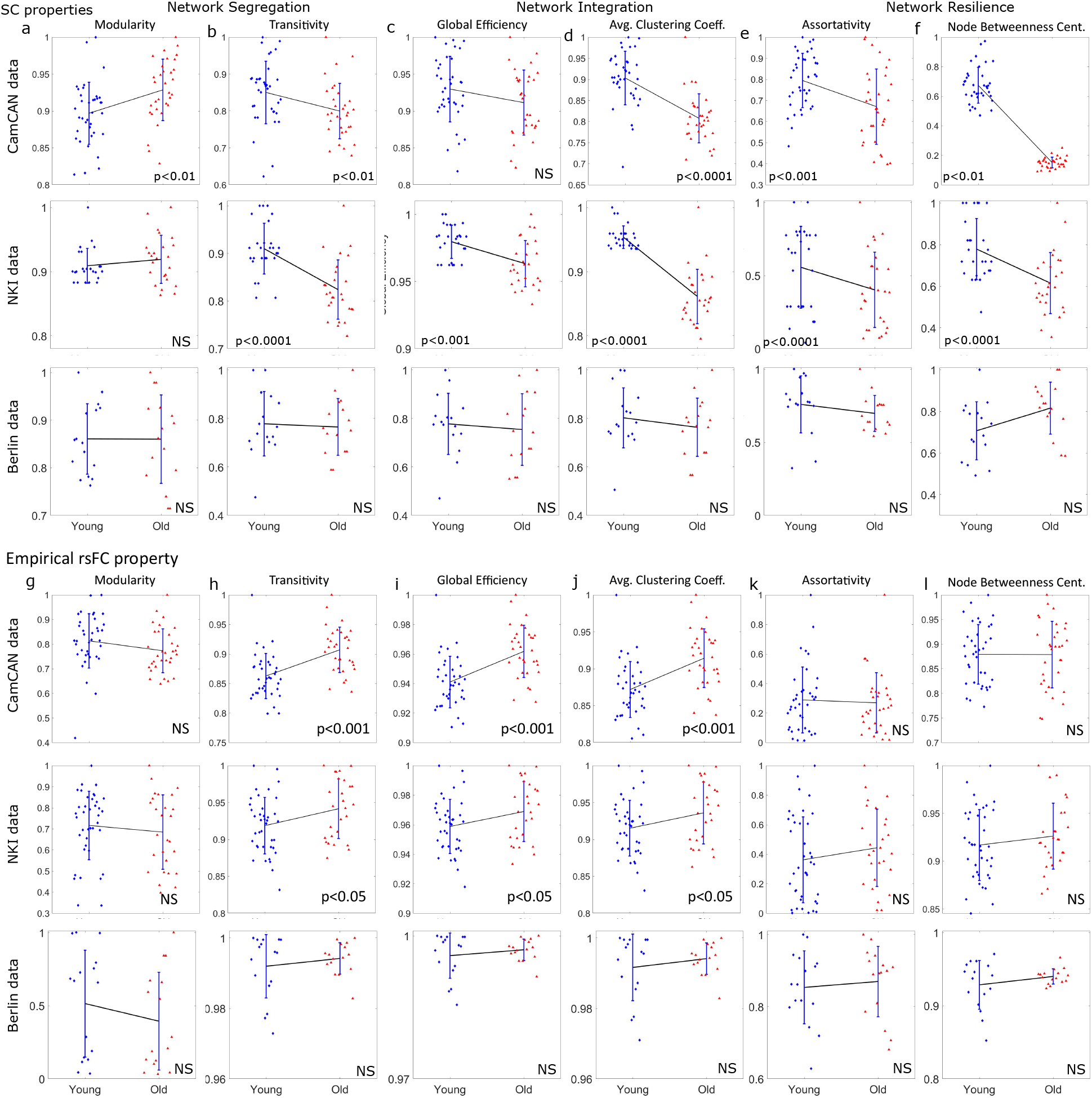
Changes in SC and functional network properties in the three data sets. We joint then by a line to show the pattern of the shift between two means of the two groups. significantly altered or unchanged network properties are determined by two-sample t-test. Number of participants: CamCAN n=37 young, n=32 older adults; NKI n=39 young, n=27 older adults; Berlin n=25 young, n=14 older adults.

### 2.2 Effect of threshold on empirical rsFC graph properties

The effect of threshold, used to generate binarized FCs, is shown in the Fig. S2. We performed test-retest validation for two different thresholds (corresponds to *p <* 0.04, 0.03 of the correlation matrix, i.e., abs(FC) ≥0.085,0.091), which are able to reproduce similar pattern of changes in the graph metrics in the three datasets.

**Figure S2.**
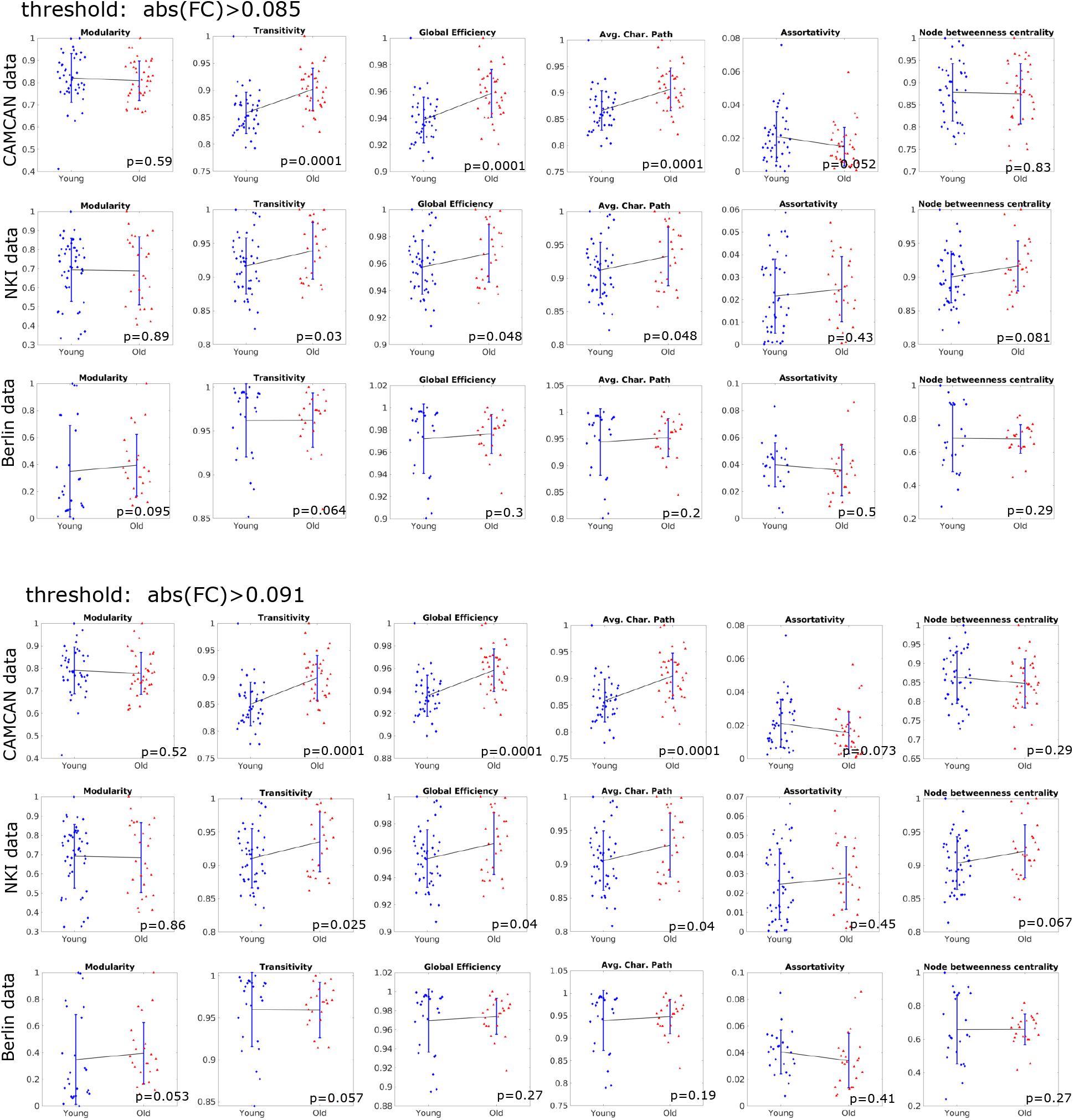
Effect of threshold to binarize the empirical FCs (Pearson correlation coefficient). Patterns of changes in the rsFC network properties in the three datasets remains stable between two age-groups for a wide range of the threshold. Number of participants: CamCAN n=37 young and n=32 older adults; NKI n=39 young and n=27 older adults; Berlin n=25 young and n=14 older adults.

### 2.3 Consistency of empirical rsFC graph properties

To test the thresholding effects on the FC network property, we defined consistency solely based on our observations on the obtained FC properties for CamCAN and NKI datasets, such that the modularity, assortativity, node-betweenness-centrality are non-significant, whereas, transitivity, global efficiency and average characteristic path are significant when comparing between the two age groups; if any of these conditions are violated we consider as not consistent. We have plotted the graph theoretical metrics (averaged across subjects for each group) against the varying threshold, shown in Fig. S3(a). The three cyan color lines in Fig. S3(a) are the threshold values (0.077, 0.085, and 0.091) used for the empirical rsFC analysis. These three thresholds correspond to the pair-wise p-values (*p<*0.05, 0.04, and 0.03) obtained during the Pearson correlation coefficient calculation applied to create the FCs from the resting state BOLD signals. Further, to assess the effect of threshold, we calculate the sparsity and percentage of remaining edges after FC binarization (*FC*^*bin*^), shown in Fig. S3(b). To evaluate sparsity, we introduce the following measure,

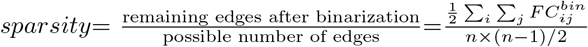

where, n = total number of brain areas. We also calculate, % of preserved edges = sparsity × 100 %, and edge losses as, loss = [1 − sparsity] × 100%. The p-values of individual metrics are unified to check the consistency of our observations on FC with varying thresholds. We separated significant and non-significant p-values as: (i) p-values of the metrics showed significant alteration between the two aging cohorts: *p*_*sig*_ = (*p*val_T_ + *p*val_GE_ + *p*val_CP_)*/*3, where, *p*val_T_, *p*val_GE_ and *p*val_CP_ represent the p-values computed for the measures transitivity, global efficiency and characteristic path length respectively when comparing between young and old. (ii) p-values of the metrics showed non-significant alteration between the two groups: *p*_*nonsig*_ = (*p*val_M_+*p*val_A_+*p*val_BC_))*/*3, where, *p*val_M_, *p*val_A_ and *p*val_BC_ represent the p-values computed for the measures modularity, assortativity and betweenness centrality respectively when comparing between young and older adults. We unified the two outcomes into a single measure, to check the consistency of our observations and amount of edge loss against threshold as, Consistency = *a*Θ(*δ* − *p*_*sig*_)+(1−*a*)Θ(*p*_*nonsig*_−*δ*), where, Θ(.) is the Heaviside function, where *δ*=0.05 and *a*=0.5. For visualization purpose, we have inverted the values, showed in Fig. S3(b) in solid blue line. The value becomes 1, only if all the observations are consistent, otherwise 0. Our results are consistent for the loss*<* 30% edges, where connected component (averaged across subjects) of the graph remains one, as shown in dashed black line. The connected component essentially means every node of a network is accessible at least via one path from any other remaining nodes. We have presented the results choosing the threshold from the shaded range, where the amount of preserved edge is 95% to 84%. The doted-red line and solid yellow lines show preserved edge and loss, respectively.

**Figure S3.**
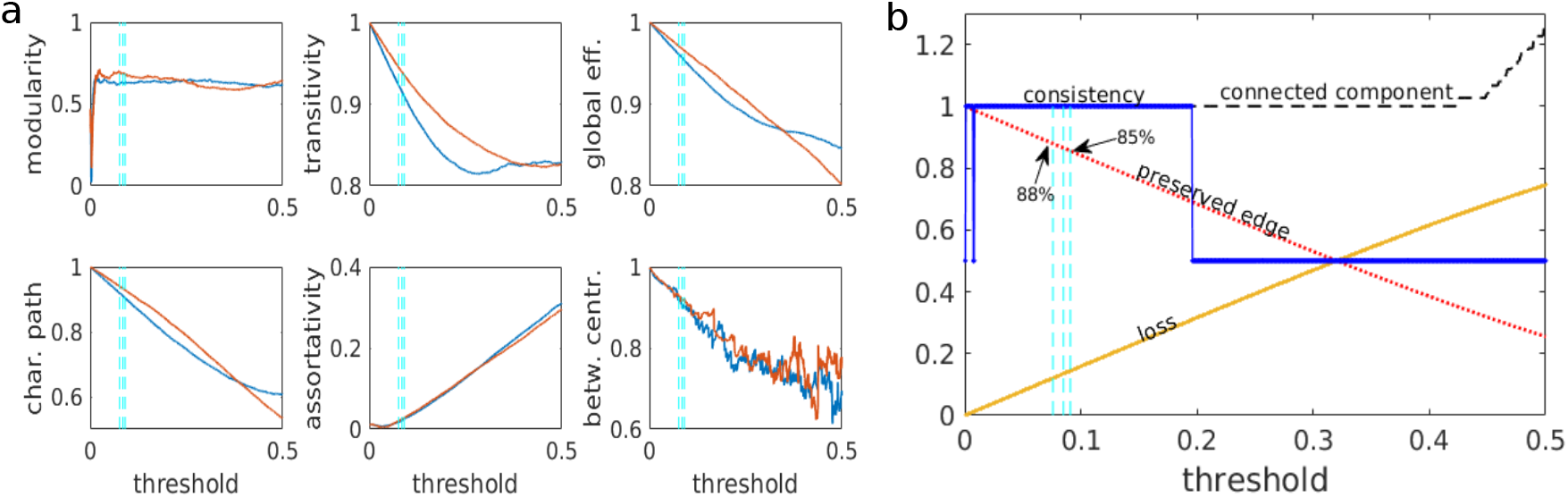
Effects of thresholding on our observations of FC properties are shown for CamCAN data. (a) Graph metrics are shown for varying threshold for young (blue) and old (red). The three cyan color lines are the threshold values ((0.077, 0.085 and 0.091) used to create the binarized empirical rsFC in our analysis. (b) Effects of the thresholding on our observations are measured by consistency, connected component, percentage of edges preserved or lost.

### 2.4 Empirical BOLD activity at resting state and tasks for CamCAN data

#### 2.4.1 Data sources and participants

The data were collected as part of stage 2 of the Cambridge Centre for Aging and Neuroscience (CamCAN) project (available at http://www.mrc-cbu. cam. ac.uk/datasets/camcan) (Taylor et al (2017); Shafto et al (2014). The fMRI data from resting state [rest] and task periods (naturalistic movie watching [mov] and sensorimotor task [SMT]) were used in the present study.

#### 2.4.2 Data preprocessing

The fMRI data for each functional run (resting state, movie watching, and SMT) were unwarped using fieldmap images, realigned to correct for motion and, slicetime corrected. EPI data were coregistered to the T1 image, transformed to montreal neurological institute (MNI) space using the warps and affine transformation from structural image (estimated using DARTEL). For region of interest (ROI) analysis, mean regional BOLD time series were estimated in 116 parcellated brain areas of Anatomical Automatic Labelling atlas (AAL) Tzourio-Mazoyer et al (2002) (available at http://www.gin.cnrs.fr/tools/aal). Preprocessed data were provided by CamCAN research consortium. Detailed overview of preprocessing pipeline can be found in Taylor et al (2017). We divided the whole data of 645 participants into two cohorts, young adults with an age range 18-32 years (mean age=26.53 ± 4.09 years), and old adults with an age range 60-88 years (mean age=72.69 ± 7.55 years). Each participant’s BOLD time series in the resting state, naturalistic movie watching, and SMTs were utilized to measure the metastability.

#### 2.4.3 Results on empirical metastability in rest and task

Patterns of brain activity appear to be more stable during cognitive operations requiring explicit attention Chen et al (2015); Cohen (2018); Elton and Gao (2015); Hutchison and Morton (2015). Existing theories suggest that spontaneous neural dynamics represent a repository of functional states from which more stable global brain states are constructed during task-based reasoning Alderson et al (2020). Therefore, metastability should be maximal when subjects are at ‘rest’ and minimal during cognitive demand Alderson et al (2020). We tested the hypothesis that large-scale brain dynamics is more metastable at rest than during the execution of an explicit task. The degree of metastability is shown in Fig. S4 for three conditions, rest, movie watching (mov) and sensory motor task (smt) by considering all participant together (left panel), young cohort (second panel), elderly subjects (third panel), and compare between younger and elderly adults (fourth panel).

**Figure S4.**
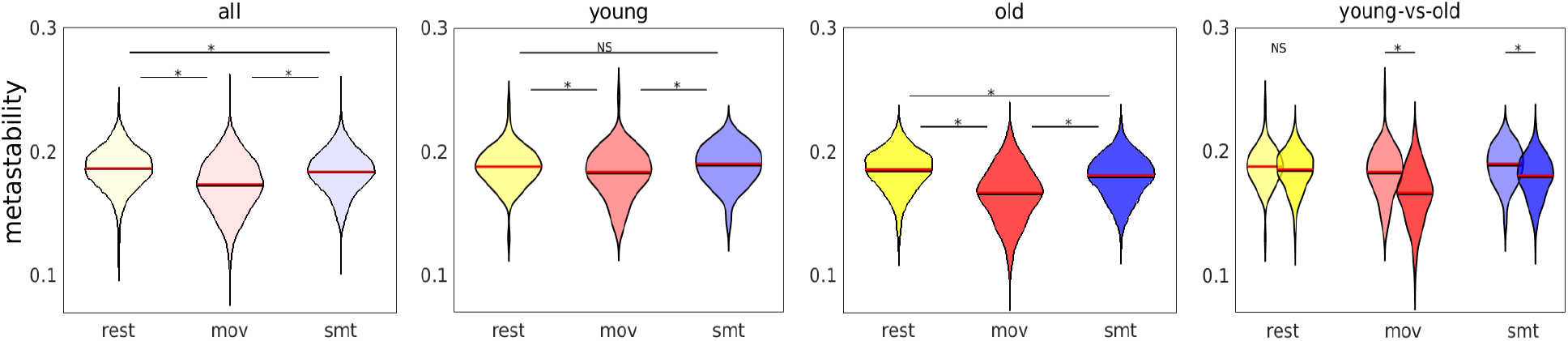
We perform independent t-test to compare empirical metastability for conditions, resting state, movie watching and sensory motor task (n=645 participants). *∗* represents *p<*0.01.

To examine any computational artifacts in our results, we tested our observations against surrogated data using phase randomization, shown in Fig. S5. Lower value of Lin’s concordance suggest that the observations from empirical data are not reproducible through any arbitrary or random process.

**Figure S5.**
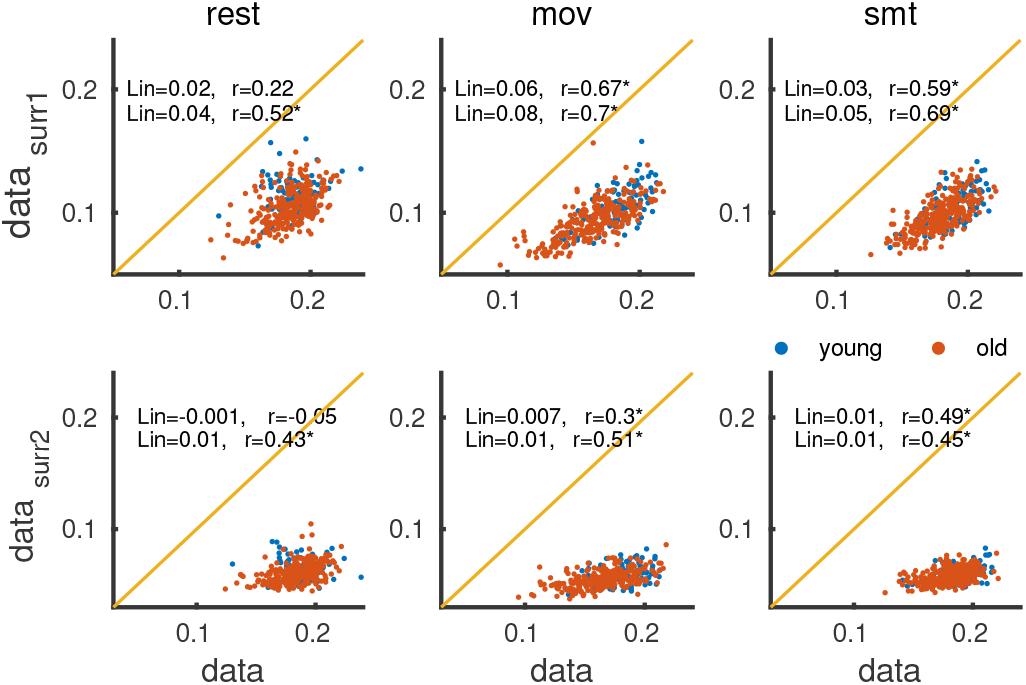
Reproducibility of the empirical metastability from surrogated data (phase randomized BOLD signal). These scatter plots depict the relationships between surrogated and empirical data metastability by Lin’s concordance for the CamCAN dataset. Lower values of Lin’s concordance suggest that random process is unable to reproduce the empirical metastability in three conditions (resting state, movie watching and sensory motor task) in the two aging cohorts. The texts show Lin’s concordance (Lin) and Pearson correlation (r); upper and lower texts for young and older cohort, respectively. *∗* indicates *p<*0.01.

## 3 Results from the Multiscale Dynamic Mean Field model

### 3.1 Multiscale Dynamic Mean Field (MDMF) model

The whole-brain dynamics is described by a set of coupled nonlinear stochastic differential equations, the current-based MDMF model Naskar et al (2021) as,

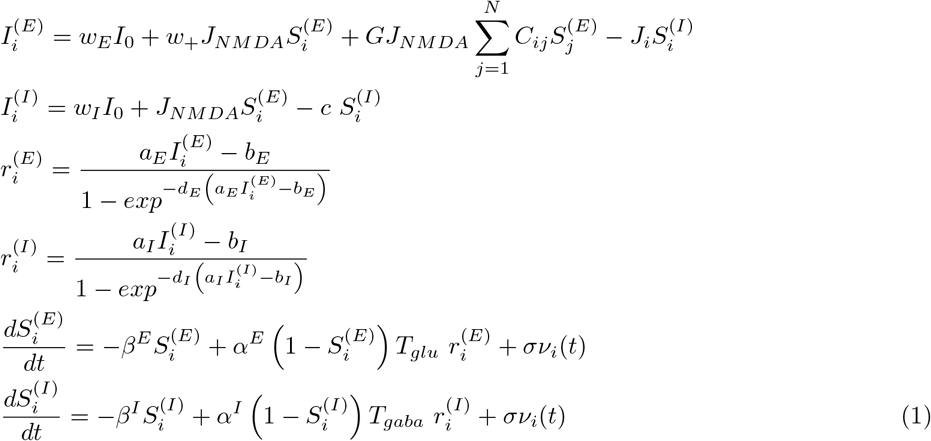

where, the subscripts *i, j* indicate brain areas, and superscripts *E, I* represent excitatory or inhibitory populations, respectively. Total number of considered brain areas is *N*, where its value is different for different parcellation choice, e.g., *N* = 68 and 188, respectively for Desikan-Killiany parcellation Desikan et al (2006) and Craddock 200 atlas Craddock et al (2012). 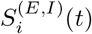 denotes the average excitatory or inhibitory synaptic gating variables at *i*^*th*^ local brain area. The time-dependent gating variables are drawn by averaging the fraction of open channels of neurons. The stochasticity is introduced by adding uncorrelated white Gaussian noise *ν*_*i*_ within the equation of two gating variables with intensity *σ* for each brain region. Further details of the model formulation and descriptions are available in the article by A. Naskar et al. Naskar et al (2021). *J*_*i*_ denotes the local synaptic coupling strength from inhibitory to excitatory. The temporal dynamics of the inhibitory feedback are given by,

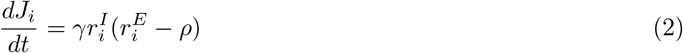

At the mean-field level, the biological complexity involved in the balance of dynamics between excitatory and inhibitory fields can be captured grossly using the mathematical implementation of the inhibitory plasticity rule Hellyer et al (2016). An inhibitory plasticity rule represents changes in *J*_*i*_(*t*) (synaptic weight) to ensure that the inhibitory current clamps to an excitatory population, maintaining homeostasis. Homeostasis is achieved with *J*_*i*_(*t*) dynamics such that the firing rate of the excitatory population is maintained at the target firing rate *ρ* = 3*Hz*, and *γ* is the learning rate in *sec*. The chosen target firing rate is the firing rate observed when the inhibitory and excitatory currents are matched.

**Table S2.** Default parameter values, and descriptions shown with references.

| Parameter | Value/Unit | Description | Ref. |
| --- | --- | --- | --- |
| $I_0$ | 0.382 nA | overall effective external input current | Deco et al (2014) |
| $w_+$ | 1.4 | local excitatory recurrence | Deco et al (2014) |
| $J_{NM DA}$ | 0.15 nA | excitatory synaptic coupling strength | Deco et al (2014) |
| $\sigma$ | 0.001 | noise intensity | Naskar et al (2021) |
| $w_E$ | 1 | scaling parameter for excitatory populations | Deco et al (2014) |
| $w_I$ | 0.7 | scaling parameter for inhibitory populations | Deco et al (2014) |
| $\alpha_E$ | 0.072 ms<sup>-1</sup>mMol<sup>-1</sup> | forward rate constants of excitatory pools | Destexhe et al (1994) |
| $\alpha_I$ | 0.53 ms<sup>-1</sup>mMol<sup>-1</sup> | forward rate constants of inhibitory pools | Destexhe et al (1994) |
| $\beta_E$ | 0.0066 ms<sup>-1</sup> | backward rate constants of excitatory pools | Destexhe et al (1994) |
| $\beta_I$ | 0.18 ms<sup>-1</sup> | backward rate constants of inhibitory pools | Destexhe et al (1994) |
| $a_E$ | 310 nC<sup>-1</sup> | parameter for input-output function | Deco et al (2014) |
| $a_I$ | 615 nC<sup>-1</sup> | parameter for input-output function | Deco et al (2014) |
| $b_E$ | 125 Hz | parameter for input-output function | Deco et al (2014) |
| $b_I$ | 177 Hz | parameter for input-output function | Deco et al (2014) |
| $d_E$ | 0.16 s | parameter for input-output function | Deco et al (2014) |
| $d_I$ | 0.087 s | parameter for input-output function | Deco et al (2014) |
| $\gamma$ | 1 C | learning rate | Naskar et al (2021) |
| $\rho$ | 3 Hz | target firing rate | Deco et al (2014) |
| $G$ | 0.5 | global coupling strength | |
| $c$ | 1 nA | local inhibitory to inhibitory strength | |
| $T_{glu}$ | 0.1-15 mMol | glutamate concentration | Naskar et al (2021) |
| $T_{gaba}$ | 0.1-15 mMol | GABA concentration | Naskar et al (2021) |

### 3.2 Model generated resting-state FC

Here we discussed how we estimate model generated functional connectivity matrix, which will be used to check the model’s predictability under certain assumptions. The simulated neural fluctuations were given by the firing rate 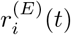 for *i*^*th*^ brain area fluctuates around a fixed value. The neural activity from the MDMF model was converted to the BOLD activity using a hemodynamic Ballon-Windkassel model Friston et al (2003). All the biophysical parameters are taken from Friston et al (2003). The simulated BOLD activity was filtered by a band pass filter and detrending operating converting into normal distribution, while removing trends and temporal mean set to zero for individual brain areas. Pearson correlation between all pair of brain regions generates the resting state model FC.

### 3.3 Synthetic glutamate-GABA estimation using MDMF model inversion

Figure S6 describes the MDMF model simulation and four steps to estimate model-based concentrations of the two neurotransmitters for a given structural connectivity of individual participants.

Step 1, in Figs. S6(a-d), involves synthetic generation of neural activity and the fitting method that allows us to project the model-generated low frequency (0.004−0.1Hz) BOLD signals and their spatiotemporal harmony, particularly fluctuations in the cortical coherence. The spontaneous brain activity at rest is captured by the empirical BOLD signals derived from resting state fMRI data. The two BOLD signals (empirical and simulated) depict hemodynamic responses of different brain areas, where pair-wise spatial correlation is the weights in the functional connectivity (FC) matrix. Detailed steps in the first stage analysis are: Fig. S6(a) We use anatomical connectivity derived from diffusion tensor imaging from the subjects of different ages. Fig. S6(b) A mean-field neuronal model is put on top of the structural connectivity matrix. Thus, the nodal dynamics is governed by the MDMF model Naskar et al (2021) and spatially connected via SC matrix. The structural connectivity describes the inter-areal interaction strength to exchange excitability across the cortical regions. We generate a model simulated neural activity, firing rate of excitatory/inhibitory populations of individual regions. The Balloon-Windkessel algorithm Friston et al (2003) then estimates the hemodynamic activity at rest. A Pearson correlation is derived from the estimated hemodynamic activity between each pair of brain regions. Thus, we approximate the functional connectivity (FC), which we mentioned as the model based or simulated FC. Figure S6(c) shows an exemplary resting-state fMRI activation map, which is used to extract empirical resting-state BOLD activity of individual subjects using CONN toolbox. In Fig, S6(d), we plot the empirical rsFC from the BOLD activity, which is used in fitting against resting-state model FC. In Step 2, the fitting measures Euclidean distance between empirical and model FCs, called mean square root error between two functional connectivity (mseFC), shown in Fig. S6(e). Simultaneously, the fluctuation in global coherence is measured by metastability, when tuning the two intrinsic parameters (*T*_*glu*_ and *T*_*gaba*_), see Fig. S6(f). We have varied the two local parameters, *T*_*glu*_ and *T*_*gaba*_ homogeneously across the whole brain and store the two measures for each pair of parameter values in the two-parameter phase diagrams, see Figs. S6(e) and S6(f).

**Figure S6.**
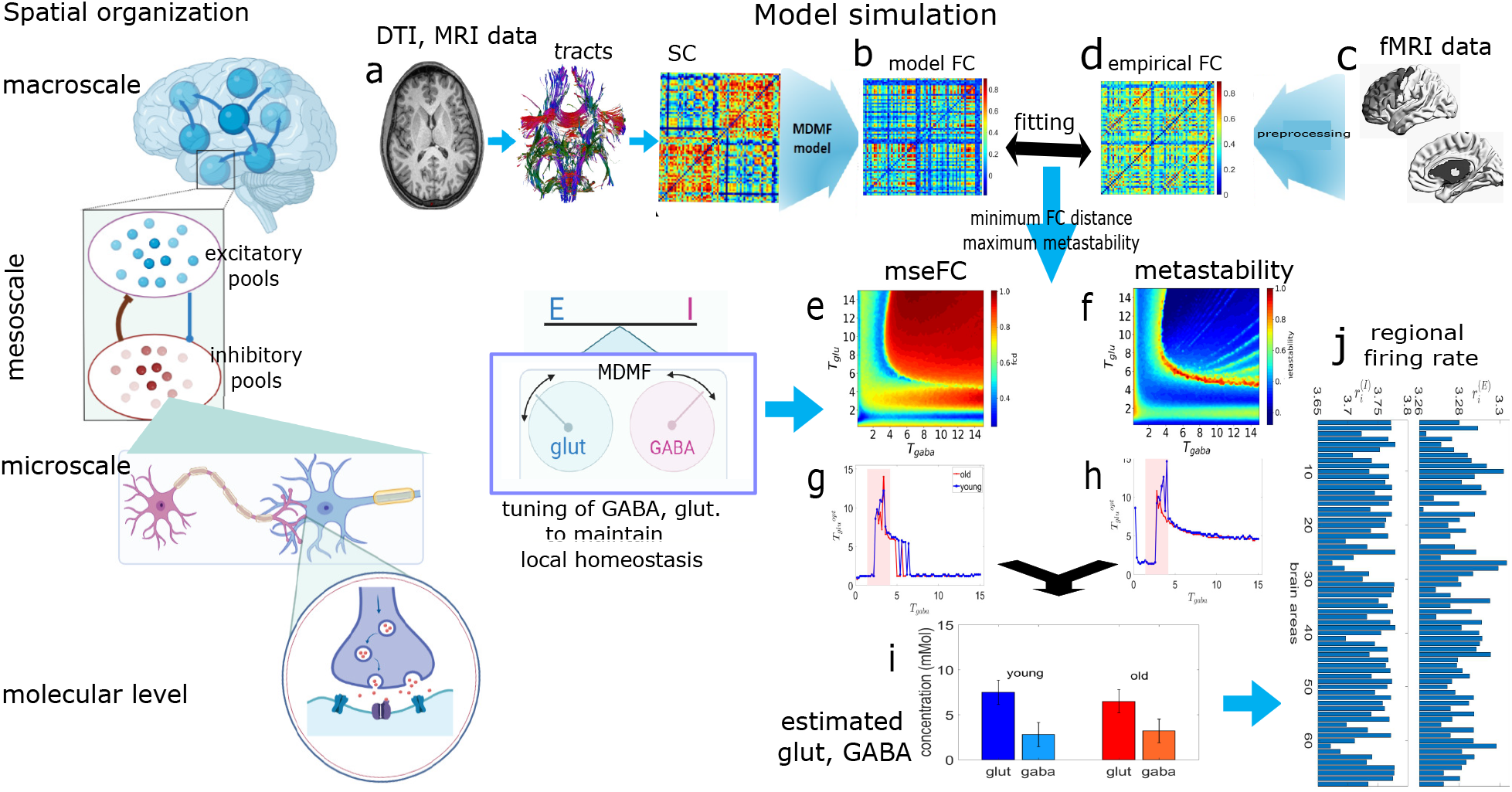
Estimation of synthetic GABA and glutamate using MDMF model. Four steps of the model simulation and synthetic result generation are shown here. (a-d) Step 1, we compare the model FC from spatiotemporal patterns to those observed in the empirical FC obtained from fMRI BOLD activity at rest. (e, f) In step 2, mseFC and metastability are stored for each pair of GABA-glutamate values. (g, h) Step 3, we estimate optimal glutamate for different values of GABA imposing the two optimal conditions. (i) Step 4, an optimal GABA is estimated from the result obtained in the previous step. Thus, an optimal set of GGC is estimated for an individual subject, based on mseFC and metastability measures. Few icons and assets are taken from the biorender (Created in https://BioRender.com).

In Step 3, see Figs. S6(g, h), we scan along the x-axis, i.e., across GABA values, and estimate optimal glutamate concentration (*T*_*glu*_) in *mMol* imposing two optimal conditions, minimum FC distance (i.e., maximum correlation between the model-generated FC and empirical rsFC), and maximum metastability, which decides optimal brain functionality. We average over the estimated glutamate values obtained from the two measures from Figs. S6(g, h).

Finally, in Step 4, GABA concentration is estimated for the homeostasis range Naskar et al (2021), indicated by shaded boxes in Figs S6(g, h) under the same optimal conditions. Thus, subject-wise optimal parameter set of *T*_*gaba*_ and *T*_*glu*_, is predicted investigating whole-brain neuroimage data (MRI, fMRI), shown using a representative result in Fig. S6(i). The firing rate remains at the desired set point of ∼4Hz, as shown in Fig. S6(j), only for the optimal choice of the neurotransmitter levels.

Further, we monitor GABA-glutamate concentrations (GGC) and their ratio (GGR) for two aging cohorts at the level of single subjects. We separate healthy subjects into two discrete age groups, young and old. Then, a possible alteration (or invariance) pattern in the two metabolites is captured over the lifespan, which has helped draw a plausible conclusion.

### 3.4 Model performance evaluation: Effects of thresholds on simulated FC graph properties

We perform test-retest validation to examine the effect of the threshold used to binarize the simulated FCs, generated considering optimal GABA-glutamate values, presented in Fig. S7. We present the results obtained for two different thresholds, which are able to reproduce a similar pattern of changes in the graph metrics in the two datasets. We refer to Fig. S1 and Fig. S2 for empirical rsFC properties to compare the pattern of age-related changes in the graph metrics against the results obtained in simulated FCs, shown in Fig. S7. The node betweenness centrality is not fully predicted by the model, though other network properties have behaved similarly to those of empirical data. Due to fewer participants in the Berlin dataset in two groups, we did not perform the training and testing simulations to assess the network properties.

**Figure S7.**
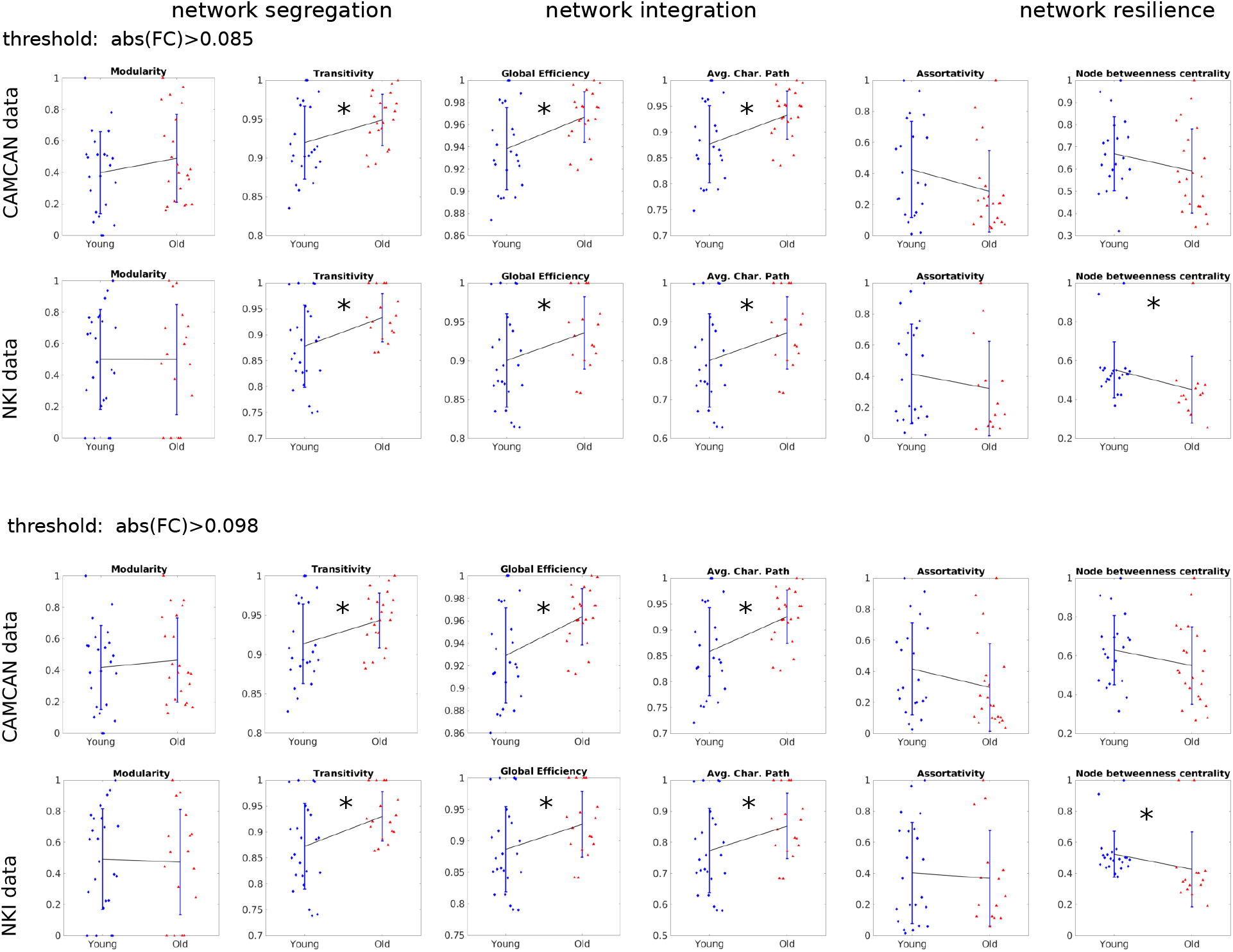
Effect of threshold to binarize the simulated FCs. The pattern of changes between young and old cohorts in the model-based FCs properties remains almost similar to that of empirical rsFC properties for a wide range of the threshold. Only node betweenness centrality metric is not fully predicted in the test cohorts; while comparing between the age groups, the statistical significance does match to the empirical observation. *∗* represents *p <* 0.05, otherwise not significant.

### 3.5 Effect of coupling strength on the GABA/glutamate estimation

Figure S8 shows the fitting for coupling strength in CamCAN data in the top row. We use FC-FC correlation, distance between empirical and simulated FCs, and FC dynamics (FCD) distance, as well. The estimated parameters are presented for various coupling strengths in the second row, whereas corresponding firing rates are presented in the third and fourth rows. The simulated FCs for both young and older adults are generated based on the estimated GABA/glutamate values, shown in the last two rows in Fig. S8. Within the shaded region, we have observed a similar pattern of Glutamate/GABA changes between the two age groups.

**Figure S8.**
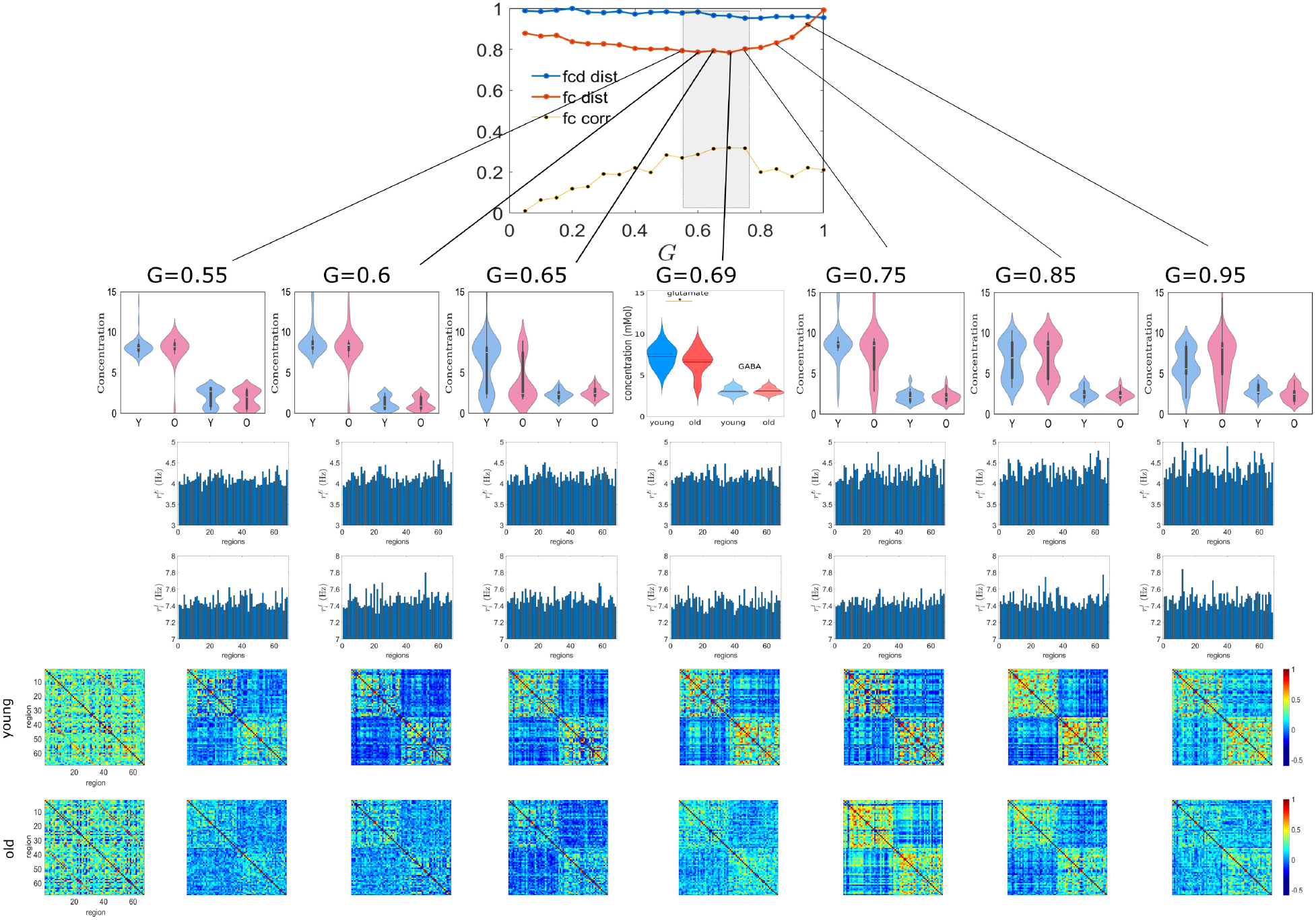
Effect of coupling strength (*G*) on parameter estimation and firing rates. The results are produced in CamCAN data. The top panel shows fitting of the coupling strength based on the distance and correlation between static FCs, and also dynamic FC distance. We have selected *G*=0.69 for our analysis in CamCAN data (young n=37, old n=32). Second row shows optimal GABA/Glutamate estimated at different *G* values. In the shaded range, we observe a similar pattern of age-related neurotransmitter changes. Excitatory and inhibitory firing rates obtained for a single participant are plotted in the third and fourth rows, respectively. The fifth and last rows present empirical and simulated FCs from one young adult and elderly participant, generated for different *G* values. Simulated FC is not the best fit and excitatory firing activity fluctuates at much higher rate when *G* is higher than optimal value.

